# Highly efficient multiplex editing: One-shot generation of 8x *Nicotiana benthamiana* and 12x Arabidopsis mutants

**DOI:** 10.1101/2020.03.31.018671

**Authors:** Johannes Stuttmann, Karen Barthel, Patrick Martin, Jana Ordon, Jessica L. Erickson, Rosalie Herr, Filiz Ferik, Carola Kretschmer, Thomas Berner, Jens Keilwagen, Sylvestre Marillonnet, Ulla Bonas

## Abstract

Genome editing by RNA-guided nucleases, such as *Sp*Cas9, has been used in numerous different plant species. However, to what extent multiple independent loci can be targeted simultaneously by multiplexing has not been well-documented. Here, we developed a toolkit, based on a highly intron-optimized zCas9i gene, which allows assembly of nuclease constructs expressing up to 32 sgRNAs. We used this toolkit to explore the limits of multiplexing in two major model species, and report on isolation of transgene-free octuple *Nicotiana benthamiana* and duodecuple (12x) *Arabidopsis thaliana* mutant lines in a single generation (T_1_ and T_2_, respectively). We developed novel counter-selection markers for *N. benthamiana*, most importantly *Sl*-FAST2, comparable to the well-established Arabidopsis seed fluorescence marker, and FCY-UPP, based on production of toxic 5-fluorouracyl in presence of a precursor. Targeting eight genes with an array of nine different sgRNAs and relying on FCY-UPP for selection of non-transgenic T_1_, we identified *N. benthamiana* mutant lines with astonishingly high efficiencies: All analyzed plants carried mutations in all genes (~112/116 target sites edited). Furthermore, we targeted 12 genes by an array of 24 sgRNAs in *A. thaliana*. Efficiency was significantly lower in *A. thaliana*, and our results indicate Cas9 availability is the limiting factor in such higher order multiplexing applications. We identify a duodecuple mutant line by a combination of phenotypic screening and amplicon sequencing. The resources and results presented provide new perspectives for how multiplexing can be used to generate complex genotypes or to functionally interrogate groups of candidate genes.

## Introduction

In genome editing applications, multiplexing may refer to targeting two or more loci by a common/shared target site, or to using multiple programmable nucleases to target several independent sites. Due to the relative complexity of construct design, there are only few examples where zinc finger nucleases or transcription activator-like effector nucleases (TALENs) were used for multiplexing (see Armario Najera *et al.*, 2019 for a recent review). Nonetheless, an impressive 107/109 genes encoding caffeic acid O-methyltransferases could be edited using a single TALEN pair in sugarcane (Kannan *et al.*, 2018). Also, two TALEN pairs were used to simultaneously edit four α(1,3)-fucosyltransferase and β(1,2)-xylosyltransferase-coding genes to reduce protein N-glycosylation in *Nicotiana benthamiana* (Li *et al.*, 2016). This is most likely the only example of “true” multiplexing with TALENs or ZFNs, as addressing multiple targets became a lot easier with the discovery of RNA-guided nucleases (RGNs; Jinek *et al.*, 2012, Cong *et al.*, 2013, Mali *et al.*, 2013).

Although exact modes of target programming differ, the provision or expression of additional guide RNAs (gRNAs) is sufficient to direct any of the commonly used RGNs, *Sp*Cas9, *Sa*Cas9 or Cas12, to multiple targets for multiplexing. For plant genome editing, the components encoding the RGN/gRNA system are most commonly stably integrated into the plant genome *via* Agrobacterium-mediated transformation. The nuclease system is subsequently expressed in plant cells, and desired mutations/edits can be separated from the nuclease-encoding transgene in subsequent generations by segregation. The majority of genome editing in different plant species was conducted using *Sp*Cas9 in combination with single guide RNAs (sgRNAs), which combine the crRNA and tracrRNA in a single molecule (Jinek *et al.*, 2012). So far, sgRNAs in T-DNA constructs for multiplexing have been expressed as individual transcriptional units (TUs) using RNA-Polymerase III-transcribed promoters (e.g., U6 or U3 promoters), or as polycistronic transcripts by RNA-Polymerase II or III promoters. In the latter case, the primary transcript requires processing, which is achieved by the endogenous t-RNA processing system, the endonuclease Csy4 or ribozyme sequences (Gao and Zhao, 2014, Nissim *et al.*, 2014, Xie *et al.*, 2015, Cermak *et al.*, 2017). An individual sgRNA unit measures 200-300 nucleotides (nt) when using Pol-III promoters (depending on regulatory elements), and can be as small as ~130 nt when using the Csy4 system. Thus, from an engineering perspective, there is little limitation for the integration of multiple sgRNA units into a plant transformation construct.

Accordingly, RGNs were frequently used in the multiplexing mode (reviewed in Armario Najera *et al.*, 2019), and comprehensive toolkits for assembly of respective nuclease-coding constructs are available (e.g., Xing *et al.*, 2014, Lowder *et al.*, 2015, Cermak *et al.*, 2017, Wu *et al.*, 2018, Hahn *et al.*, 2019). However, reduced efficiencies can be expected when an increasing number of sgRNAs is used for multiplexing, as individual sgRNAs will compete for the common nuclease core. Also, it is a common concern that repetitive sequences of sgRNA-coding blocks may lead to silencing of transgene expression *in planta*, and direct repeats, also in sgRNA arrays, were reported as prone to recombination (e.g., Vidigal and Ventura, 2015, Ding *et al.*, 2019, and references therein). These are most likely the reasons there are less examples where increased numbers of sgRNAs (≥ 6) were used for multiplexing. In rice, a system in which RGNs appear to be particularly effective, seven out of eight targeted *FT-like* genes were mutated using eight individual Pol-III driven sgRNA units (Ma *et al.*, 2015). In another study, a tRNA-gRNA array containing eight gRNA units was used to mutate four rice *MPK* genes (Minkenberg *et al.*, 2017). In dicots, multiplexing with six or more sgRNAs was used, among other species, in *Nicotiana benthamiana*, *N. tabaccum*, *Solanum lycopersicum, S. pimpinellifolium* and *Arabidopsis thaliana* (reviewed in Armario Najera *et al.*, 2019). E.g., arrays of six and eight sgRNAs were used for editing domestication-associated genes in *S. pimpinellifolium* (Zsogon *et al.*, 2018), and seven sgRNAs were used for editing of *N. benthamiana* genes required for N-glycosylation (Jansing *et al.*, 2019). In both studies, not all targets could be mutated at once. In *A. thaliana*, six sgRNAs were used to target six ABA receptor-encoding genes (Zhang *et al.*, 2015). In this study, a sextuple mutant line could be identified in the T_2_ generation, but was not separated from the transgene, although this is critical for unequivocal discrimination of germline-transmitted and somatic mutations. Overall, it thus remains unclear how many target sites can be addressed simultaneously and efficiently by multiplexing.

In this study, we explore the efficiency and limits of multiplex-editing in the two major dicot model species *A. thaliana* and *N. benthamiana*. We first extended a previous toolkit for assembly of constructs with up to 32 sgRNAs, and also tuned the toolkit for improved efficiencies by incorporation of a highly intron-optimized Cas9 gene, zCas9i (Grützner *et al.*, 2020). For multiplexing in *N. benthamiana*, we developed new markers for transgene counter-selection, and target eight genes by an array of nine different sgRNAs. In *A. thaliana*, we target 12 different genes with 24 sgRNAs. We show that very high efficiencies can be obtained by multiplexing in *N. benthamiana.* We identify biallelic mutations at almost all target sites in randomly selected, non-transgenic T_1_ individuals, and observe an unexpected over-representation of homozygous mutations. We also isolate a transgene-free duodecuple mutant line from editing with 24 sgRNAs in Arabidopsis, but lower efficiencies are observed. Our results show that Cas9 availability becomes the limiting factor in higher order multiplexing applications, while recombination or silencing of transgenes do not appear problematic. The high multiplex editing efficiencies we report open up new perspectives for the generation of complex genotypes and for functional analysis of numerous candidate genes by RGNs.

## Results

### Adaptation of a vector system for high efficiency multiplexing

We aimed to explore the limits and efficiency of multiplex genome editing in two major model species, *A. thaliana* and *N. benthamiana*. Therefore, we first adapted our previously developed Dicot Genome Editing (pDGE) vector system (Ordon *et al.*, 2017) for improved efficiency, streamlined selection procedures and higher-order multiplexing with up to 32 sgRNAs (Figure 1). The pDGE vector system consists of “shuttle” vectors for preparation of sgRNA transcriptional units (TUs) and preassembled “recipient” vectors (Figure 1a,b). Recipient plasmids contain a *ccdB* cassette (Figure 1a), which can be excised by *Bsa*I/*Eco*31I and replaced by sgRNA TUs in a GoldenGate cloning reaction. For enhanced efficiency, recipient vectors were equipped with a highly intron-optimized Cas9 gene, zCas9i (Grützner *et al.*, 2020), either under control of an Arabidopsis *RIBOSOMAL PROTEIN S5a* (RPS5a) promoter fragment (Tsutsui and Higashiyama, 2017, Ordon *et al.*, 2019) or a 35S promoter fragment. Furthermore, recipient plasmids contain a positive selection marker (resistance to glufosinate (BASTA), kanamycin or hygromycin) and additional markers for positive and/or negative selection. These markers allow the rapid identification of non-transgenic individuals from segregating populations, and were established previously (FAST marker for Arabidopsis, Shimada *et al.*, 2010), or will be described in later paragraphs.

**Figure 1:**
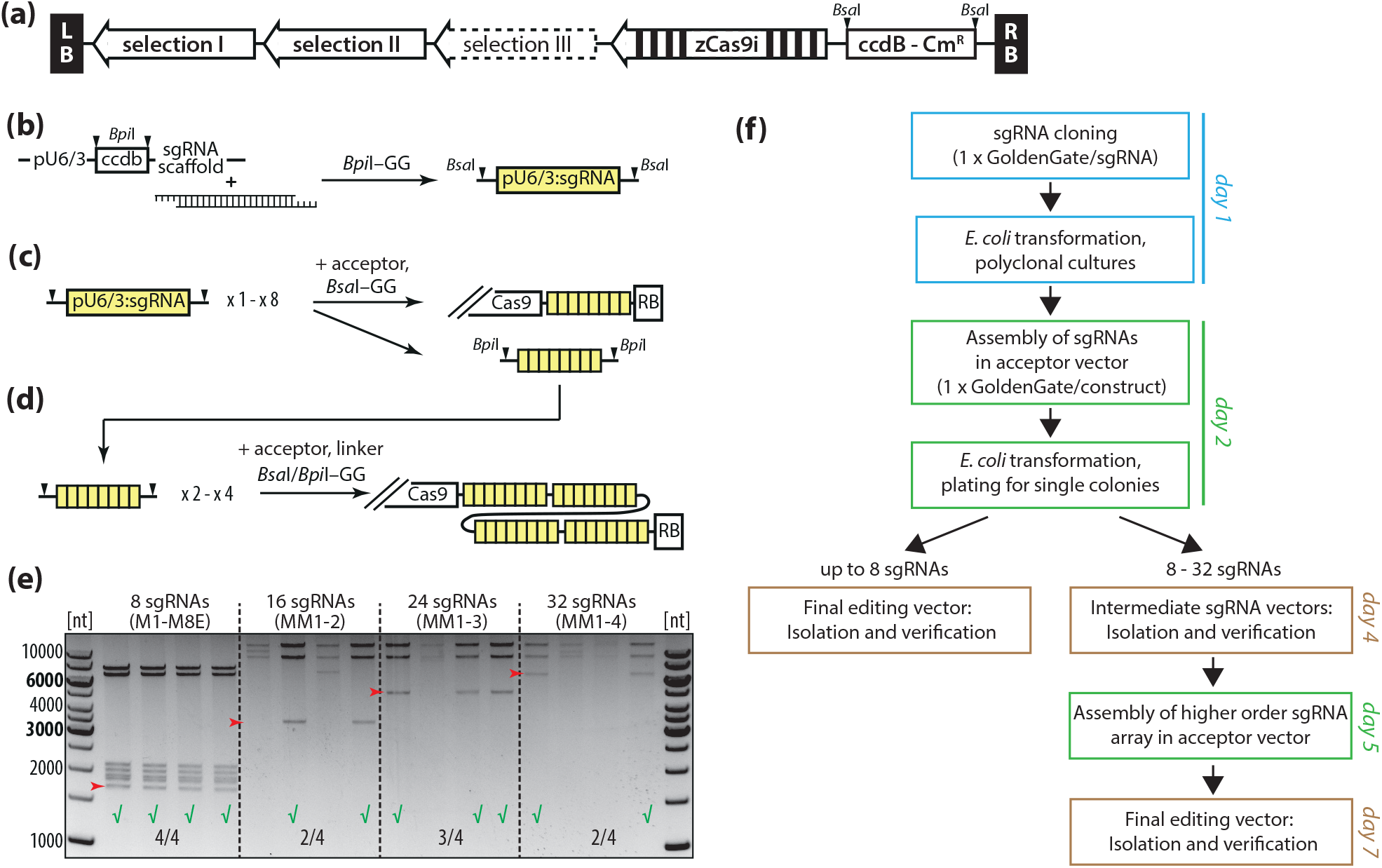
Adaptation of a vector system for enhanced efficiency and multiplexing with up to 32 sgRNAs. **(a)** General scheme of pDGE recipient/nuclease vectors. Empty nuclease vectors used in this work contained up to three selection markers for positive/negative selection in Arabidopsis or *N. benthamiana*. A highly intron-optimized Cas9 gene, zCas9i (coding for Cas9 with two nuclear localization signals), was incorportated for improved editing efficiencies (Grützner et al., 2020). A *ccdB* cassette can be excised by *Bsa*I, and replaced by an sgRNA array *via* GoldenGate cloning. **(b)** Generation of sgRNA transcriptional units (TUs) by cloning of hybridized oligos into shuttle vectors. Shuttle vectors contain either an *At*U6-26 promoter element, or U3/U6 promoter elements from tomato. Shuttle vectors are available with different *Bsa*I overhangs, determining their position in an sgRNA array. Presence of *ccdB* in empty vectors allows polyclonal processing of the GoldenGate reaction (Ordon et al., 2017). **(c)** Assembly of sgRNA arrays. Up to eight sgRNA TUs may be mobilized from loaded shuttle vectors *via* a *Bsa*I GoldenGate reaction directly into nuclease vectors, or into intermediate „multi-multi” vectors. In the latter, sgRNA arrays are flanked by *Bpi*I restriction sites for subsequent higher level assembly reactions. **(d)** Assembly of up to 32 sgRNA TUs in a nuclease vector. sgRNA arrays are mobilized from intermediate multi-multi vectors into nuclease vectors *via* a *Bsa*I/*Bpi*I GoldenGate reaction. Use of respective end-linkers allows assembly of 2, 3 or 4 arrays into any nuclease vector. **(e)** Efficiency of cloning reactions. Four clones each from assembly of 8, 16, 24 or 32 sgRNAs in a nuclease vector were randomly selected for plasmid isolation, and DNA was digested with *Pst*I/*Hind*III (8 sgRNAs; direct assembly into a nuclease vector) or *Hind*III (higher level assemblies employing multi-multi vectors). The band corresponding to the respective sgRNA array is marked with a red arrowhead. **(f)** General timeline for assembly of nuclease vectors. Editing constructs containing up to 8 sgRNA TUs can be obtained within 4 days. Including an additional day for sequence-verification of intermediate sgRNA array constructs, nuclease vectors with up to 32 sgRNAs can be obtained within 7 days.

sgRNAs TUs are prepared in shuttle vectors by cloning of hybridized oligonucleotides *via* a *Bpi*I/*Bbs*I GoldenGate reaction (Figure 1b). In shuttle vectors containing an *At*U6-26 promoter fragment, we exchanged the sgRNA scaffold (Dang *et al.*, 2015) and modified an overhang that was identified as sub-optimal using web tools (www.tools.neb.com; Pryor *et al.*, 2020). We also generated additional shuttle vectors containing either U3 or U6 promoter fragments from tomato (*S. lycopersicum; Sl*U6/U3). When tested in transient efficiency assays, the *Sl*U6/U3 promoter elements resulted in similar nuclease activity as the previously used *At*U6-26 promoter fragment (Figure S1). The *Sl*U6/U3 shuttle vectors can thus be considered an alternative to shuttle vectors containing the *At*U6 promoter fragment.

Mobilized from shuttle vectors, sgRNAs assemble into arrays of two, four, six or eight TUs, directly in recipient vectors (Figure 1c). We prepared a set of intermediate “multi-multi” cloning vectors to enable for assembly of arrays >8 sgRNA TUs. Following the same principle as the assembly of sgRNA arrays in recipients, intermediate cloning vectors can also harbor up to eight sgRNA TUs, and were prepared for four different positions in a final sgRNA array. Thus, up to 32 sgRNA TUs can be assembled into a recipient vector in a GoldenGate reaction employing *Bsa*I and *Bpi*I and a respective end-linker (Figure 1d; Weber *et al.*, 2011). We tested the efficiency of assembling 8, 16, 24 or 32 sgRNAs in a recipient vector (Figure 1e): Although the efficiency of higher-order assemblies using multi-multi vectors was reduced in comparison to direct assembly of 8 sgRNAs (theoretically, a more demanding assembly involving more DNA inserts), ≥ 50 % of tested clones were positive. Overall, this extended pDGE system allows for simple and high-fidelity assembly of nuclease constructs containing up to 8 sgRNAs within 4 days, or containing up to 32 sgRNAs within 6-7 days (Figure 1f).

The incorporation of multiple selection markers and the intron-optimized zCas9i into pDGE vectors leads to relatively large T-DNA regions of final plant transformation constructs (e.g., ~ 20 kb and ~ 17 kb in vectors used for editing in *N. benthamiana* and *A. thaliana* in experiments described in following sections). It is a common concern that larger T-DNAs show reduced transformation efficiencies. However, we consistently observed high transformation efficiencies with our constructs in Arabidopsis and *N. benthamiana* (Figure S2), and it should be noted that natural T-DNAs are commonly 10 – 30 kb in size (Gelvin, 2003).

### Development of negative selection markers for isolation of transgene-free genome edited N. benthamiana plants

Upon continuous presence of a nuclease-coding transgene, a given genotype may not remain stable in subsequent generations or may vary between different parts, organs or tissues of a plant. It is therefore imperative to use non-transgenic individuals from multiplex editing for faithful genotyping and phenotypic analyses. In Arabidopsis, the FAST marker provides a convenient method not only for positive transgene selection *via* seed fluorescence (Shimada *et al.*, 2010), but also for negative selection (Castel *et al.*, 2018, Ordon *et al.*, 2019). For *N. benthamiana*, a comparable marker system has not been developed. This may be considered even more critical, as *N. benthamiana* transformants often contain more than one T-DNA insertion, at least under our conditions. As a consequence, large populations have to be screened to identify non-transgenic segregants. We therefore set out to develop alternative negative selection markers for use in *N. benthamiana*.

In consecutive editing experiments and transformations, we tested in total eight different cassettes as potential negative selection markers in *N. benthamiana* (Figure 2a). We initially tested the FAST marker, as was used in Arabidopsis, but this did not result in detectable seed fluorescence. We then tested a fusion of the Arabidopsis 2S3 promoter (Kroj *et al.*, 2003) with genes coding for mCherry, tagRFP or 2xtagRFP. These cassettes were combined with the pepper (*Capsicum annuum*) *Bs3* gene under control of its own promoter in transformation constructs. The *Ca-Bs3* gene is normally not expressed, but provokes cell death when induced by the transcription activator-like effector AvrBs3 (Römer *et al.*, 2007, Boch *et al.*, 2014). We aimed to use inducible cell death as a marker.

**Figure 2:**
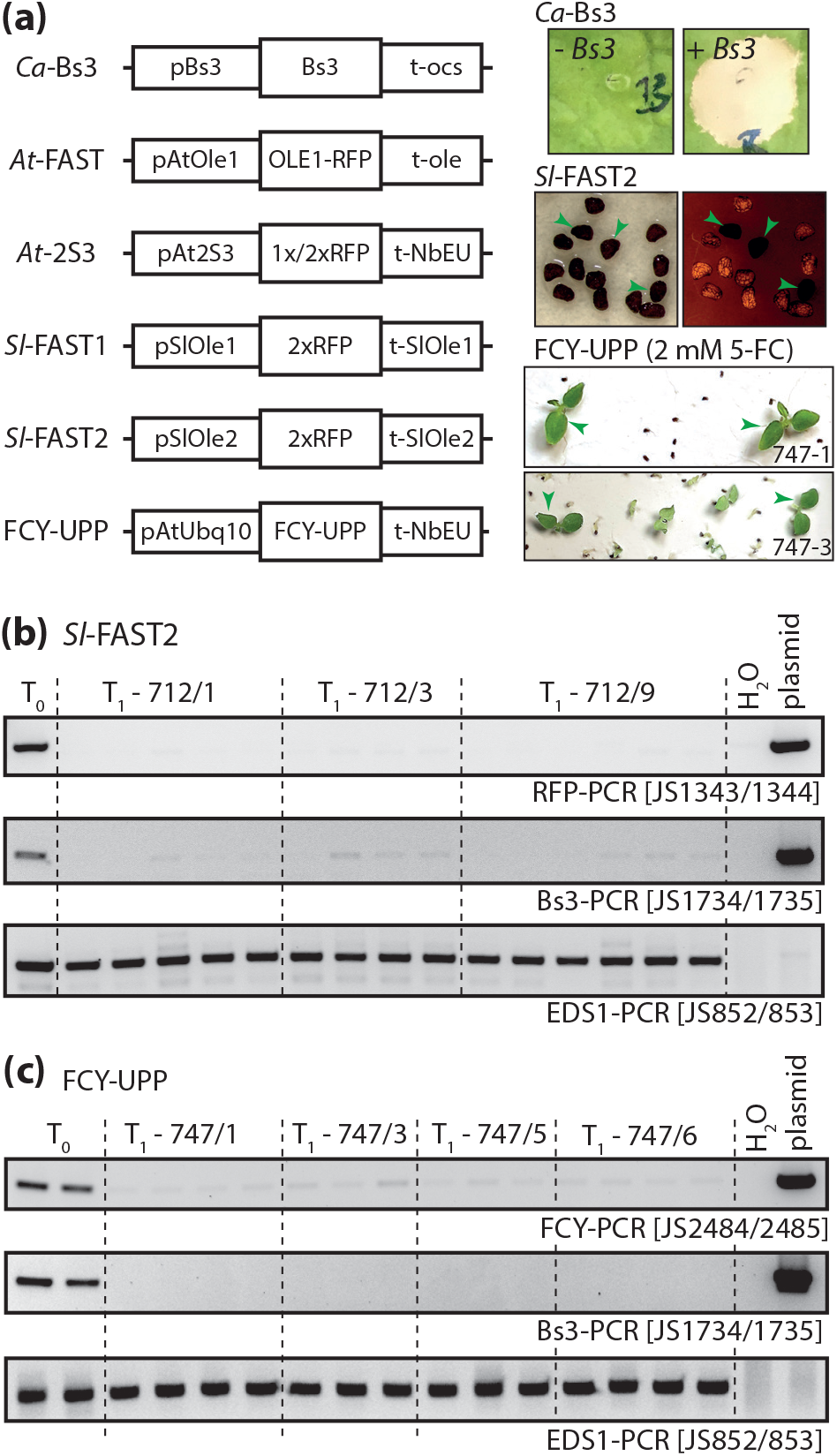
Markers for selection of non-transgenic *N. benthamiana* from segregating T_1_ populations. **(a)** Overview on tested marker cassettes. Representative pictures of markers with high predictive power are shown. To query for presence/absence of *Bs3*, approximately 4 weeks-old plants were infiltrated with a *Pseudomonas fluorescens* strain for translocation of AvrBs3 (Gantner et al., 2018). Phenotypes were recorded 3 dpi, images show a *Bs3*-transgenic plant in comparison to wild-type control. For the *Sl*-FAST2 marker, seeds were briefly imbibed by placing them on wet Whatman paper, and imaged using a stereo microscope under white light conditions (left image) or under UV illumination using an RFP filter (right image). Arrowheads mark non-fluorescent seeds. For the FCY-UPP marker, T_1_ seeds were sown on quartz sand plates supplemented with 1/4 MS solution containing 2 mM 5-fluorocytosine (5-FC). Pictures were taken after 12 days. Arrowheads mark plants representative for segregants used for PCR-genotyping. **(b)** Evaluation of counter-selection using the *Sl*-FAST2 marker. Non-fluorescent seeds as shown in (a) were selected and germinated on soil. Leaf tissues were sampled 14 days after germination and used for DNA extraction and PCR-genotyping with the indicated primer pairs. The *EDS1* amplicon was included as a control for DNA quality. The *Bs3* and *RFP* amplicons query presence/absence of the T-DNA. **(c)** Evaluation of counter-selection using the FCY-UPP marker. Plants were grown on quartz sand plates containing 2 mM 5-FC as shown in a). Leaf tissues of healthy plants were sampled 12 days after germination and used for DNA extraction and PCR-genotyping with the indicated primer pairs. Amplicons as in b), but a primer pair specific for *FCY-UPP* was used in place of the *RFP* amplicon.

The 2S3:mCherry marker cassette resulted in weak seed fluorescence which was not visible by direct observation through the eyepiece of a stereo microscope, but only upon documentation of seeds using a digital camera and extended exposure times (5 -10 sec.; Figure S3). Fusing either one or tandem copies of tagRFP-T-coding genes with the 2S3 promoter and use of a strong terminator (*N. benthamiana* extensin; Diamos and Mason, 2018) only marginally improved seed fluorescence. The 2S3:mCherry/RFP cassettes allowed selection of non-transgenic plants with an acceptable predictive power (Figure S3): Without any pre-selection, most plants (20/22) tested positive for the transgene. After selection by seed fluorescence, the transgene was not detected in plants from 3/5 independent T_1_ families. However, counter-selection was not successful for the remaining two families we tested (Figure S3), and the selection procedure including imaging of seeds was not straightforward. Thus, the 2S3 promoter fragment has only limited utility for counter-selection in *N. benthamiana*. By contrast, cell death induction upon expression (by Agrobacterium) or translocation (by *P. fluorescens*) of the AvrBs3 TALE perfectly coincided with the presence of the transgene (Figure S3c). *Ca-Bs3* proved highly reliable as a marker under our conditions, and was included in all further transformation constructs.

We further sought to use promoter and terminator fragments from oleosin-coding genes of a solanaceous plant to adapt the FAST system for use in *N. benthamiana* (Shimada *et al.*, 2011). We identified two oleosin-coding genes in the tomato genome, and assembled two additional markers, *Sl*-FAST1/2, using respective promoter and terminator fragments (Figure 2a). Fluorescence of seeds of *Sl*-FAST1-transgenic *N. benthamiana* was comparable to that previously obtained with the 2S3 promoter fragment. However, seeds of plants containing the *Sl*-FAST2 marker demonstrated strong seed fluorescence that was, although weaker than the FAST marker in Arabidopsis, easily discernable by direct observation through the eyepiece of a stereomicroscope (Figure 2a). It should be noted that fluorescence was not visible with dry seeds, but appeared within seconds when seeds were imbibed, e.g. by placing them on a wet piece of Whatman paper for observation. When the *Sl*-FAST2 marker was used for seed pre-selection, none of the plants grown from selected, non-fluorescent seeds tested positive for the transgene (Figure 2b). Hence, *Sl*-FAST2 is a useful dominant and non-invasive marker for *N. benthamiana*.

Last, we tested a fusion of a yeast (*Saccharomyces cerevisiae*) cytosine deaminase-coding gene (*ScFCY*) and an *E. coli* phosphoribosyl transferase-coding gene (*EcUPP*) under control of an Arabidopsis *Ubiquitin10* promoter fragment as negative selection marker (Figure 2a). FCY converts 5-fluorocytosine (5-FC) into the antipyrimidine 5-fluorouracil (5-FU), which blocks thymidine synthetic processes and is incorporated into DNA and RNA (Longley *et al.*, 2003). *Ec*UPP enhances the RNA incorporation pathway and thus the toxicity of 5-FU (Tiraby *et al.*, 1998). Plants lack cytosine deaminase activity. Therefore, 5-FC only becomes toxic to plants in presence of the *FCY-UPP* gene; a similar fusion was recently used for tissue-specific genetic ablation (Leonhardt *et al.*, 2020). We wanted to avoid working under sterile conditions and therefore first tested toxicity of 5-FC on quartz sand plates supplemented with MS solution (Davis *et al.*, 2009) using a previously reported *FCY-UPP*-transgenic Arabidopsis line (under 35S promoter control; Leonhardt *et al.*, 2020). 5-FC did not have any adverse effects on the growth of wild-type plants, but became toxic to the transgenic line at concentrations of 1mM or higher (Figure S4). We then sowed wild-type *N. benthamiana* and T_1_ seeds from transformation of a construct containing the FCY-UPP marker on quartz sand plates containing 1 or 2 mM 5-FC (Figure S5). As with Arabidopsis, we did not observe adverse effects of 5-FC on growth of wild-type *N. benthamiana*. Strikingly, from the two independent T_1_ families from transformation of the *FCY-UPP* construct that were initially tested, many seeds failed to germinate for one, while most seeds germinated for the other line, but only few plants developed normally (Figure S5). We next tested four independent T_1_s in parallel (including the two tested previously) using 2 mM 5-FC. For three families, we observed again that many seeds failed to germinate (Figure 2a, line 747-1), while most seeds germinated for the last line, but only few plants (3/~150) developed normally (Figure 2a, line 747-3). We took tissue samples as early as 12 days after sowing and genotyped plants by PCR (Figure 2c). Indeed, none of the plants selected *via* the FCY-UPP system tested positive for the transgene. This suggests that, e.g., for line 747-3, we most likely identified the only three non-transgenic plants from ~ 150 seeds basically without any effort and in a single step. Thus, the *FCY-UPP* marker appears extremely reliable and convenient as a negative selection marker, at least in *N. benthamiana*.

### One-shot generation of octuple mutant N. benthamiana plants in a single generation

Having identified suitable counter-selection markers for *N. benthamiana*, we next wanted to test multiplex editing in this system. We decided to target eight genes by a construct encoding nine different sgRNAs (Figure 3a,b; note that sgRNAs 5/9 in the array were inadvertently chosen identical). Due to its allotetraploid nature, the genome of *N. benthamiana* often contains two highly similar gene copies. When possible, we designed sgRNAs to target both copies. Using this strategy and the complement of nine different sgRNAs, each gene (except one) contained two target sites, giving a total of 15 target sites. The sgRNA array was assembled in three steps using the position 1 and position 2 intermediate cloning vectors in pDGE792, a recipient containing zCas9i under control of the Arabidopsis RPS5a promoter, the Bar selection cassette for positive selection, and the *Ca-Bs3* and *FCY-UPP* cassettes for negative selection. We selected non-transgenic plants by insensitivity to 5-FC on quartz sand plates and confirmed absence of the transgene by PCR. Four DNAs, originating from two sister plants each from two independent T_1_ families, were used to amplify sgRNA target sites by PCR. For one locus, we detected a deletion, while PCR products of remaining loci had the size expected for the wild type (Figure S6). Amplicons were Sanger-sequenced to inspect for point mutations. Astonishingly, all plants had mutations in all targeted genes, most of which were biallelic, and we noted an unexpected overrepresentation of homozygous mutations (Figures 3c, S7-10). Indeed, wild-type sequences were detected only in two plants, at four target sites. In total, we scored 112/116 analyzed alleles as mutant (96,5% efficiency), although the precise sequences of alleles could not be determined in all cases. This suggests that complex higher order mutants can be generated and obtained in *N. benthamiana* with stunningly high efficiencies. Targeting eight genes with nine sgRNAs, we did apparently not reach the limits of multiplexing in this system.

**Figure 3:**
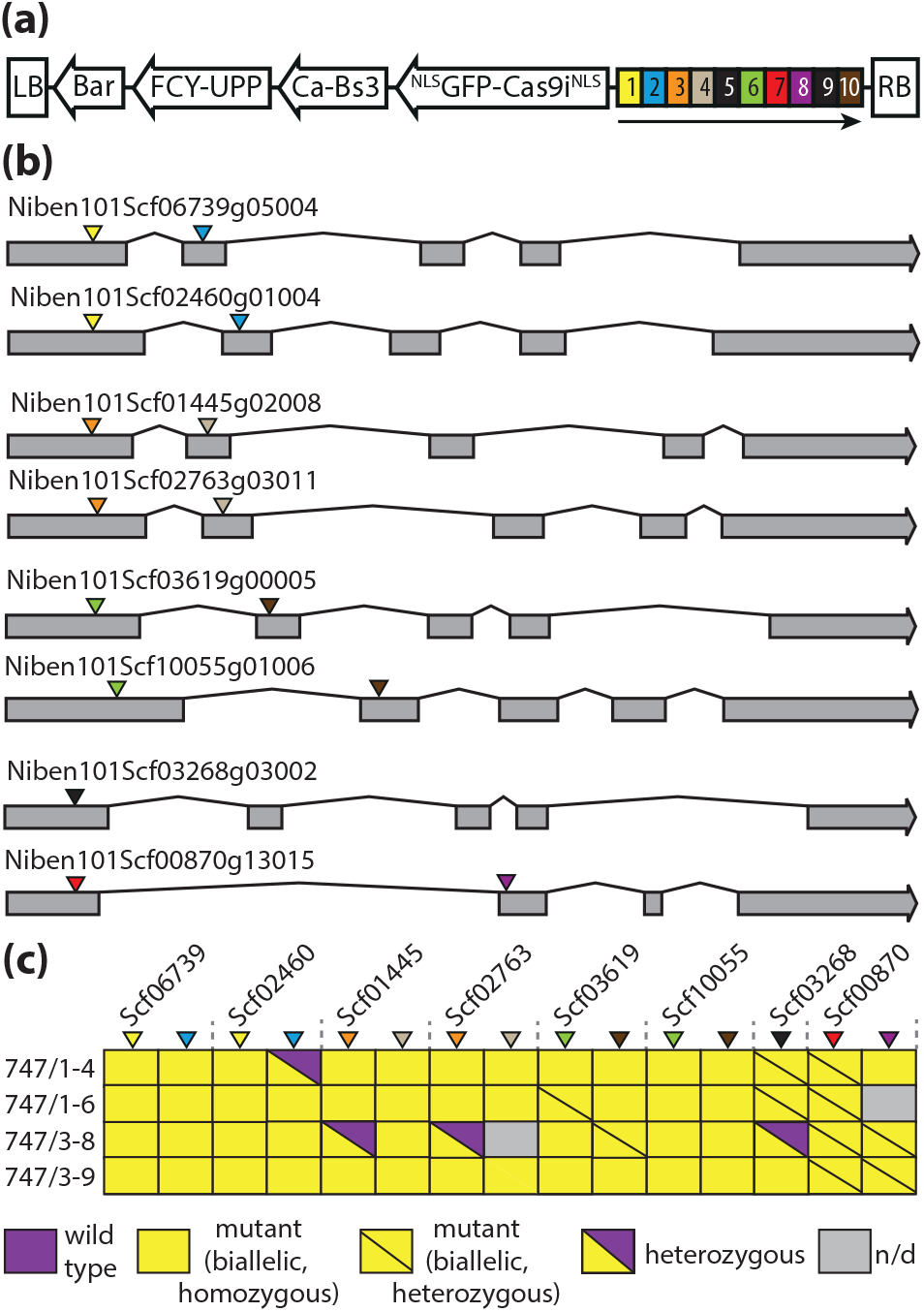
One-shot generation of octuple mutant *N. benthamiana* lines. **(a)** Schematic drawing of the construct used for plant transformation. Expression of the NLSGFP-Cas9NLS fusion is controlled by an Arabidopsis RPS5a promoter fragment and a chimeric triple terminator (35S+*Nb*ACT3+Rb7; Diamos et al., 2018). The arrow indicates the direction of transcription of the sgRNA transcription units (TUs). sgRNAs 5/9 are identical; expression of all sgRNAs is under control of an *At*U6-26 promoter element. **(b)** Overview on genes that were targeted by multiplex editing. Coloured triangles indicate sgRNA target sites; the color code corresponds to the sgRNA array shown in panel (a). **(c)** Summary of results from genotyping of four transgene-free T_1_ plants from transformation of the construct depicted in (a). T_1_ seeds were sown on quartz sand plates containing 5-fluorocytosine for selection of non-transgenic plants. After 12d, one cotelydon was cut and used for DNA extraction and PCR-genotyping. Amplicons covering the sgRNA target sites were sequenced directly (see Figures S7 - S10 for chromatograms). In total, 112/116 analyzed sgRNA target sites were scored as edited.

### Editing 12 genes by 24 sgRNAs in Arabidopsis thaliana: Analysis of primary transformants

We tested multiplexing in Arabidopsis by targeting 12 different genes by a construct containing an array of 24 sgRNA TUs (Figure 4a). Selected target genes included several nucleotide-binding domain– leucine-rich repeat-type resistance genes (e.g., *RPP2*, *RPP4*, *RPS5*) and developmental regulators (e.g., *ERECTA*, *TOO MANY MOUTHS*, *GLABROUS1*). Target genes and sites are listed in Table S1. As previously described for multiplexing in *N. benthamiana*, each gene was targeted by two different sgRNAs. A respective construct was assembled in pDGE347 (containing the FAST marker, the Bar gene and zCas9i under control of the Arabidopsis RPS5a promoter), using the position one, two and three intermediate cloning vectors. Furthermore, sgRNAs directed against *ERECTA* (*ER*) or GLABROUS1 (*GL1*) were, either in pairs or simultaneously, directly assembled in pDGE347 to generate three control constructs. Although single and double mutants could be induced with high efficiencies in the T_1_ generation in previous transformations with the intron-optimized zCas9i (Grützner *et al.*, 2020), we expected that efficiencies would decrease due to competition of many different sgRNAs for the limiting Cas9 nuclease core. Furthermore, we suspected that the 24 successive blocks of (with the exception of the 20 nt variable section of the sgRNA) identical sequence repeats within the sgRNA array might be prone to recombination events in *E. coli*, in *Agrobacterium*, during T-DNA transfer, or after T-DNA integration into the plant genome.

**Figure 4:**
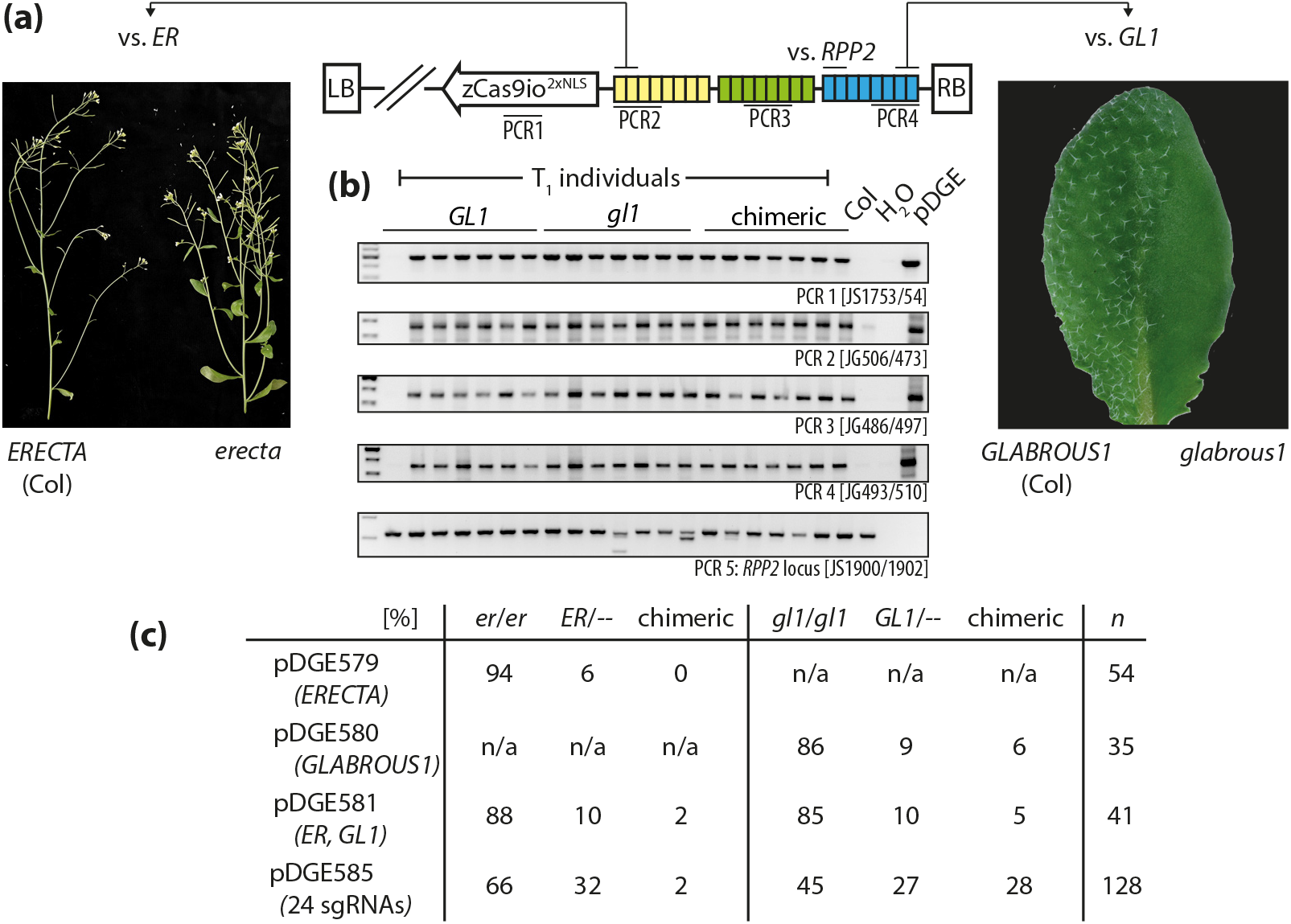
Analysis of primary (T_1_) transformants from multiplex-editing in Arabidopsis. **(a)** Schematic drawing of the plant transformation construct (pDGE585) containing 24 sgRNA transcriptional units (TUs). Blocks consisting of 8 sgRNA TUs each from intermediate cloning steps (positions 1 - 3) are depicted in yellow, green and blue, respectively. PCR amplicons used for verification of T-DNA integrity are indicated. Insets show phenotypes resulting from editing of loci targeted by sgRNAs flanking the array. **(b)** Verification of T-DNA integrity in primary transformants. Phenotypes (trichome development; *GLABROUS1*) of primary transformants (selected for resistance to BASTA) used for DNA extractions are indicated. Untransformed wild type (Col) and the transformation vector (pDGE) were included as controls. PCR 5 amplifies a fragment of the *RPP2* locus trageted by two nucleases encoded by the multiplexing constructs as shown in a). **(c)** Frequency of plants with *erecta* and *glabrous* phenotypes in T_1_ plants from transformation of indicated constructs. pDGE579 and 580 contained sgRNA TUs (two each) for targeting of *ERECTA* and *GLABROUS1*. pDGE581 contained the four sgRNA TUs from pDGE579/580 in a single construct. pDGE585 contained the four sgRNA TUs from pDGE581, and 20 additional sgRNA TUs. Values indicate the number of primary transfromants showing a respective phenotype in percent. *n* indicates the total number of primary transformants analyzed for each construct.

To assess possible recombination events, the construct containing the 24 sgRNA TUs was transformed into Columbia-0 (Col) wild-type plants. Primary transformants were selected by resistance to BASTA, and DNA extracted for PCR-genotyping (Figure 4b). Different primer pairs reveal the presence of the Cas9 gene and three different blocks each encompassing four sgRNA TUs spread along the sgRNA array (Figure 4a). Furthermore, a primer pair for an amplicon of the endogenous *RPP2a* locus, which was also targeted by paired nucleases, was included for genotyping. In total, 36 randomly chosen primary transformants were analyzed (Figures 4b; S11). One transformant did apparently not contain the T-DNA (Figure 4b, lane1). In another transformant, only two of the three fragments covering the sgRNA array could be PCR-amplified (Figure S11), suggesting a partial integration of the T-DNA or a recombination event. All remaining transformants were positive for the tested amplicons, and signals indicative of deletions at the *RPP2* locus were detected for some transformants (Figure 4b, PCR 5, lanes 11, 14; Figure S11). Thus, direct repeats of the sgRNA array did not induce frequent recombination events, and an intact T-DNA region was likely transferred to most transformants (34/36, 94 %).

The 24 sgRNA array was constructed in such way that it was flanked by sgRNAs for targeting the *ER* and *GL1* loci; the same sgRNAs that were also incorporated in control constructs. Inactivation of *ER* and *GL1* leads to altered shoot morphology and absence of trichomes, respectively (Oppenheimer *et al.*, 1991, Torii *et al.*, 1996); phenotypes that can easily be scored by visual inspection (see insets in Figure 4). We used appearance of *er* and *gl1* phenotypes to evaluate genome editing efficiency of the 24 sgRNA construct in the T_1_ generation. Control constructs induced mutations in *ER* and *GL1* at frequencies > 80 %, and the efficiency at individual loci did not change markedly when both loci were targeted simultaneously (Figure 4c). By contrast, editing efficiencies dropped by approximately 30-40% in transformants that received the 24 sgRNA construct. Also, the frequency of chimeric plants, which can easily be recognized for *gl1*, appeared to increase (Figure 4a). These observations support that the availability of Cas9 indeed becomes a limiting factor upon co-expression of numerous sgRNAs, which compete for integration into the nuclease core. Nonetheless, it should be noted that T_1_ efficiencies remained at roughly 50 %, and thus considerably high.

### Editing 12 genes by 24 sgRNAs in Arabidopsis thaliana: Analysis of T_2_ segregants, and isolation of a transgene-free duodecuple (12x) mutant

We further analyzed the occurrence of mutations from multiplexing in Arabidopsis in the T_2_ generation, with the goal of identifying a transgene-free duodecuple mutant line directly in the T_2_ generation. From T_1_ analysis, it became obvious that this would not be as easy a task as in the *N. benthamiana* multiplex editing trial (Figure 3). We therefore first analyzed functionality of six different genes (*RPP2*, *FKD1*, *RPS2*, *RPP4*, *RPS5* and *TMM*) in phenotypic assays, which involved infection assays with different strains of the plant pathogenic bacterium *Pseudomonas syringae* or the oomycete *Hyaloperonospora arabidopsidis* and microscopic analyses of destained leaves (Figure 5a). Mainly T_2_ families from primary transformants scored as *er gl1* double mutants or that had at least one of these mutations were included in analyses (see Table S2 for details). Plants were grown from seeds selected for absence of fluorescence to avoid confounding effects due to presence of the T-DNA and occurrence of novel somatic mutations. On average, 13 plants/family from 28 independent T_2_ families were analyzed for each phenotype; ~ 2500 phenotypes were scored. These analyses revealed that mutations at all six loci were present within the population, but occurred at different frequencies. E.g., the *rps2* phenotype, which was most frequent, was observed in 89 % of the families analyzed, and occurred in 60 % of T_2_ plants. The *rpp4* phenotype was present in only 29 % of families and 7 % of T_2_ plants. Different editing efficiencies were expected due to variable on-target efficacy of sgRNAs, even though we attempted to mitigate this effect by targeting each locus with two sgRNAs.

**Figure 5:**
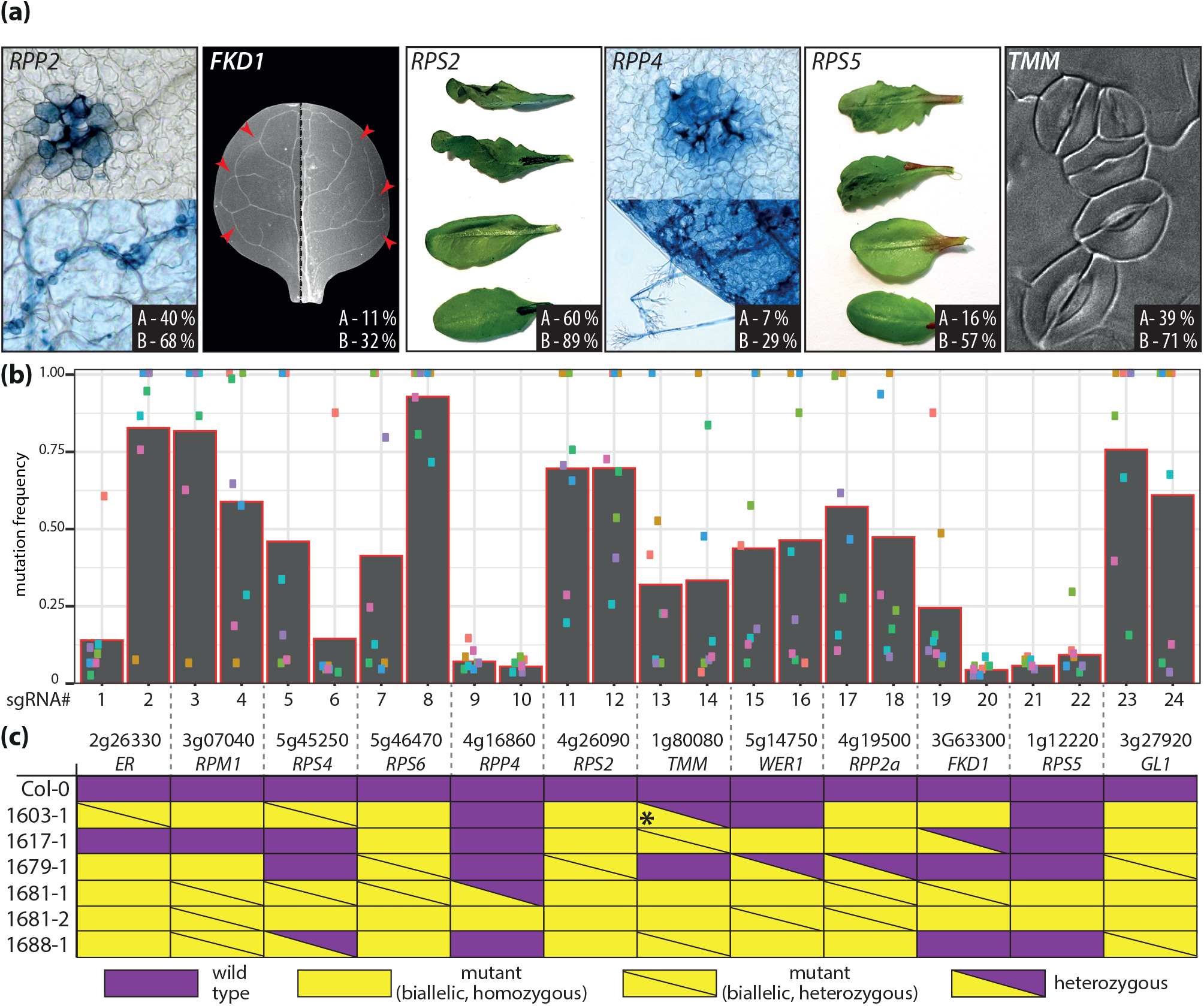
Analysis of transgene-free T_2_ segregants from multiplex-editing in Arabidopsis. **(a)** Representative images of phenotypes assessed in T_2_ segregants. The percentage of plants which were scored mutant for a respective phenotype among all tested segregants (A) and the percentage of T_2_ families, in which the phenotype was detected (B), are indicated (see Table S2 for additional details). In average, 13 segregants from 28 independent T_2_ families were analyzed for each phenotype. **(b)** Mutation frequency at individual sgRNA target sites. DNA was extracted from 10 transgene-free segregants per T_2_ family, and used as template for amplification of target loci. Pooled amplicons were sequenced by Illumina technology, and data was analyzed as described in methods to evaluate sgRNA efficiency. Mutation frequencies are color-coded for T_2_ pools (#1603; #1606; #1611; #1613; #1673; #1679; #1681, #1696). **(c)** Analysis of target locus integrity in single plants. Alleles at target loci from single transgene-free segregants were analyzed by amplicon sequencing. Each locus was targeted by two different sgRNAs, and a gene was scored as “mutant” if both detected alleles contained a mutation in at least one sgRNA target site. Details on detected alleles are provided in Figure S12. * - the mutant allele detected at the *TMM* locus in plant 1603-1 contained an in-frame deletion (6 nt), which might not disrupt TMM function.

To corroborate phenotypic analyses and to identify alleles present at target sites, we conducted short read sequencing of PCR amplicons. Prior to sequencing, absence of the T-DNA/Cas9 was verified by PCR. Ten T_2_ segregants per family were pooled and respective DNAs used for PCR amplification of target sites. PCR amplicons were then subjected to short read sequencing. Eight different families were analyzed, and the editing frequency at each target site calculated (Figure 5b). Results from amplicon sequencing revealed differences in sgRNA efficiency at target site resolution (in contrast to phenotypic analyses, which provided target locus resolution), and correlated well with our phenotypic analyses. Low mutation efficiencies were detected at both target sites within *RPP4* and *RPS5*, and respective mutant phenotypes were rare within the population (Figure 5a,b). Similarly, the *rps2* mutant phenotype was most frequent, and high mutation efficiencies were detected for both sgRNAs targeting this locus.

Next, we used short read sequencing of target loci in individual plants. Two lines were selected based on phenotypic analysis: Most plants of family #1681 appeared to be mutant at all tested loci (Table S2), and also line #1688 was highly mutagenic. Remaining lines were selected randomly. All lines we analyzed contained mutations within at least seven of the genes that were targeted by the 24 sgRNAs (Figure 5c). As observed before for sequencing of pools, most plants contained wild-type alleles at the *RPS5* and *RPP4* loci, again confirming low efficiency of respective sgRNAs. However, the *rpp4* phenotype was detected by phenotypic analyses within family #1681 (Table S2), and one of the two segregants we analyzed indeed carried disruptive mutations within all of the 12 target genes, with edits at 40/48 target sites (Figures 5c, S12). Thus, enhanced efficiencies achieved by intron-optimization of the Cas9 sequence (Grützner *et al.*, 2020) coupled with a high degree of multiplexing enabled us to isolate a duodecuple Arabidopsis mutant within a single generation.

## Discussion

It was previously estimated that at least one redundant paralog is present for more than 50 % of all Arabidopsis genes (Armisen *et al.*, 2008, Chen *et al.*, 2010), and less than 10 % of all single mutant lines from T-DNA collections were attributed a phenotype in previous systematic analyses (e.g., Cutler and McCourt, 2005, Bolle *et al.*, 2013). It is thus conceivable that, in many cases, gene function can be revealed only by analysis of higher order mutants. The availability of –omics data, allowing the selection of genes for reverse analyses e.g. based on phylogenetic analyses and/or transcriptomic data, combined with CRISPR tools for reverse genetics, now provide new opportunities to reveal these masked functions. Here, we explore the limits of multiplex gene editing in two different model systems, and generate octuple and duodecuple mutant plants in *N. benthamiana* and Arabidopsis, respectively. Our data demonstrate the efficiency of CRISPR tools, and that even large gene families or groups of genes of interest can be targeted for inactivation and gene functional analyses.

The extended and improved pDGE vector system presented here provides one route to assemble nuclease constructs for higher order multiplex-editing approaches. In its current state, the pDGE system can handle up to 32 sgRNA units, but can be extended following the same design principles. Alternatively, highly complex nuclease constructs may also be assembled using the Modular Cloning system (Weber *et al.*, 2011, Hahn *et al.*, 2019, Grützner *et al.*, 2020). Modular Cloning offers maximal flexibility, while the pDGE system, based on preassembled recipient vectors containing a range of different zCas9 expression cassettes and selection markers, provides simplicity of use. A manual for cloning of nuclease constructs with pDGE vectors also containing brief laboratory protocols for selection of non-transgenic individuals from segregating populations is provided in Appendix 1. Most plasmids of our pDGE system were submitted to Addgene upon description in a preprint (Barthel *et al.*, 2020). These are readily available as single plasmids and will also be available as a kit together with additional zCas9i plasmids (Grützner *et al.*, 2020). Additional data on functionality of plasmids is provided in Figure S1. Further plasmids not included in this deposit but described here (e.g., pDGE687 containing the *Sl*-FAST2 and pDGE792 the FCY-UPP marker) will be made available.

### Transgene counter-selection strategies

In multiplex plant genome editing applications, the identification of a genotype or a phenotype of interest may eventually involve the screening of large numbers of plants. Only transgene-free individuals allow a truly meaningful association of genotype and phenotype. In Arabidopsis, established selection systems based on seed fluorescence provide means for convenient and simple transgene counter-selection (Stuitje *et al.*, 2003, Bensmihen *et al.*, 2004, Shimada *et al.*, 2010). In our experiments, a handful of T_2_ plants counter-selected by the FAST marker tested positive for the transgene in later molecular analyses. We consider mis-phenotyping on the basis of seed fluorescence as unlikely, and assign these rather to accidental carry-over of fluorescence-positive seeds or additional, partial insertions not tagged by FAST. In contrast to Arabidopsis, negative selection markers for *N. benthamiana* were, as far as we are aware, not previously described.

Here we tested different cassettes for their suitability for counter-selection (Figure 2a). We identified the *Ca-Bs3* gene as an extremely reliable, simple and fast marker for *N. benthamiana*. E.g., we recently screened a population of > 200 plants T_1_ plants from a line that contained several independent transgene insertions. We identified three segregants non-responsive to AvrBs3 by infiltration (with *P. fluorescens* AvrBs3, < 1h), which were subsequently confirmed as non-transgenic. Another benefit of *Ca-Bs3* as marker is that it can also be used to confirm presence of the transgene in primary transformants, as soon as plants recover after transfer from tissue culture to soil.

Furthermore, we established *Sl*-FAST2 as a seed fluorescence marker for *N. benthamiana* (Figure 2a,b). Since *Sl*-FAST2 is based on regulatory elements from tomato, it is plausible to assume that it will be possible to use this system also in other *Solanaceae* species. Although used for counter-selection here, *Sl*-FAST2 may also be used for positive selection, e.g. for the determination of segregation ratios, or to control homo-/heterozygosity of seed stocks.

The FCY-UPP marker is probably the most versatile counter-selection selection system we used (Figure 2a,c). The selection strategy is not new (Perera *et al.*, 1993, Stougaard, 1993, Leonhardt *et al.*, 2020 and references therein), but it had previously not been considered in genome editing applications. In our hands, the FCY-UPP marker worked flawlessly for counter-selection. In addition, cultivation under non-sterile conditions on quartz sand plates allowed screening of large numbers of plants with minimal effort and requirements (Figure 2). An added advantage of the FCY-UPP marker is that its use is facultative, since the chimeric gene is inert in the absence of 5-FC. Thus, plants may also be cultivated for another generation in presence of the transgene, potentially for accumulation and fixation of additional mutations, or primary transformants may be used in crosses to mobilize the transgene in additional backgrounds. This is not the case e.g. for a system developed in rice, in which transgene-positive seeds fail to develop (He *et al.*, 2018). The FCY-UPP system is based on the absence of cytosine deaminase activity, which is required to convert the non-toxic 5-FC to toxic 5-FU. Higher eukaryotes including plants lack cytosine deaminase activity. Accordingly, the FCY-UPP marker should be universally applicable in plant genome editing; only transcriptional control by suitable regulatory elements will be required to adapt the marker for diverse species. We therefore expect that FCY-UPP could be widely used for plant genome editing applications in the future.

### Genome editing in dicots using zCas9i controlled by the RPS5a promoter

Editing efficiencies, at least when relying on transgenic expression, ultimately depend on nuclear amounts and availability of the Cas9 nuclease core and sgRNAs in transformants, as well as their expression in germline cells (Wang *et al.*, 2015, Mao *et al.*, 2016, Ordon *et al.*, 2017, Grützner *et al.*, 2020). Characterization of the zCas9i gene (included in pDGE vectors used here) revealed that it results in higher protein accumulation in comparison to non-intron-optimized Cas9 genes, and presence of two nuclear localization signals improves nuclear import (Grützner *et al.*, 2020). For editing in Arabidopsis, we previously identified the RPS5a promoter element as highly suitable in direct comparisons (Ordon *et al.*, 2019), and it also proved very efficient together with zCas9i (Grützner *et al.*, 2020). However, so far pRPS5a-driven Cas9 was only used in Arabidopsis; mainly 35S:Cas9 was employed for editing in *N. benthamiana* and other *Solanaceae* species.

Here, we used pRPS5a:GFP-zCas9i (combined with a 35S+*Nb*ACT3+Rb7 “triple terminator”; Diamos and Mason, 2018) for editing in *N. benthamiana*, and obtained efficiencies well exceeding previous reports (Figure 3; e.g., Gantner *et al.*, 2019, Jansing *et al.*, 2019). The rationale for using pRPS5a-driven Cas9 for multiplexing was that we assumed mutagenic activity would be maintained in germline tissues of primary transformants throughout development, whereas only mutations occurring early in tissue culture-based regeneration might be germline-transmitted with p35S. Indeed, we generally recovered only one type of mutations in a respective T_1_ population when using 35S:Cas9 in previous experiments, supporting this notion. By contrast, we found different alleles among sister plants from the same T_1_ population at several instances in experiments described here (Figures S7-S10), suggesting that these mutations arose at later developmental stages. Another observation was that we recovered an unexpectedly high number of homozygous mutations in our experiments. Lacking direct comparisons of promoters for editing in *N. benthamiana*, we cannot conclude which precise element of our editing system is at the basis of the observed mutagenic activity. However, we propose that the RPS5a promoter might be particularly suitable for (multiplex) editing, as it may allow obtaining a heterologous mixture of mutant alleles from a limited number of primary transformants. This strategy may readily be transferable to other plant species.

### Multiplex-editing for generation of higher order mutants or for candidate gene interrogation

Multiplex trials reported here show that complex genotypes, such as octuple or duodecuple mutants, can be generated in a single generation by genome editing, albeit with largely differing efficiencies in the two model species we tested (Figures 3 – 5).

In *N. benthamiana*, we recorded efficiencies > 95% targeting eight genes with nine different sgRNAs, without any prior experimental validation of sgRNAs. Three out of four randomly chosen, transgene-free T_1_ segregants most likely were functionally octuple mutants (Figure 3). Thus, efficient mutagenesis can also be expected when editing with even more sgRNAs. In transient efficiency assays, we detected a decrease of Cas9 activity at an individual target site when including 23 or more, but not 15, non-targeting sgRNAs into a construct. However, stable transgenic lines will be required to determine the limits of multiplexing in *N. benthamiana*.

In Arabidopsis, efficiencies at individual loci (*ER*, *GL1*) dropped sharply when the 24 sgRNA array was used (Figure 4), and isolation of the duodecuple mutant line required extensive phenotypic pre-selection (Figure 5, Table S2). These results suggest that Cas9 availability became the limiting factor with increasing numbers of sgRNAs. There were no indications for frequent recombination of sgRNA units, which is in agreement with the notion that recombination events mainly occur upon use of lentiviral vectors, or may be selected for upon introduction of selective phenotypes (Najm *et al.*, 2018, Reis *et al.*, 2019, and references therein). Thus, alternative sgRNA expression systems are unlikely to improve overall performance when using large numbers of sgRNAs in multiplexing. However, further enhancement of nuclease activity might be obtained e.g. by co-expression of the TREX2 exonuclease (Cermak *et al.*, 2017, Weiss *et al.*, 2020), or by using a different nuclease such as *Sa*Cas9, which outperformed *Sp*Cas9 in Arabidopsis at least in some contexts (Wolter *et al.*, 2018).

It has been well-documented that mutagenic activity differs between individual primary transformants, and that editing at a primary locus increases the likelihood of editing at further loci (e.g., Bollier *et al.*, Li *et al.*, 2020). Similarly, we detected mutant phenotypes at higher frequencies in families that were already scored as *er gl1* double mutants in the T_1_ generation (Table S2). Key to isolation of a duodecuple mutant line was to identify plants mutated at *RPP4*, *RPS5* and *FKD1* loci; respective sgRNAs demonstrated low efficiencies according both to phenotypic analyses and calculation of efficiencies based on pooled amplicon sequencing (Figure 5a,b; Table S2). On the one hand, this highlights the urgent need for reliable methods for prediction and/or selection of efficient sgRNAs. On the other hand, this also points out a workflow for isolation of complex genotypes from multiplexing: In a first round of analysis, short read sequencing of pooled amplicons from multiple primary transformants, as performed here (Figure 5b), can be used to identify loci with low mutation rates. In a second round of analysis, primary transformants with high mutagenic activity can be pre-selected based on presence of mutations in one or several of these key loci. Analysis of T_2_ segregants will likely allow straightforward isolation of the desired genotype, as cleavage of targets of inefficient sgRNAs shall function as an efficient marker for co-editing of remaining sites (Li *et al.*, 2020, see also Symeonidi *et al.*, 2020).

We see two potential applications for massive multiplexing in plant gene editing applications. This study shows that higher order mutants can be generated. Furthermore, gene sets or families may be targeted by multiple sgRNAs, either by constructs of defined composition or by multiplexing libraries, to unravel overlapping or redundant functions. The newly developed selection markers extend the applications of RGNs as directed forward genetics tools *via* the analysis of transgene-free T_1_/T_2_ segregants for a phenotype of interest without prior knowledge of the genotype.

## Materials and Methods

### Plant growth conditions and transformation

*N. benthamiana* wild-type plants were cultivated in a greenhouse with a 16-h light period (sunlight and/or IP65 lamps (Philips) equipped with Agro 400 W bulbs (SON-T); 130–150 μE/m^2^*s; switchpoint;100 μE/m^2^*s), 60% relative humidity at 24/20°C (day/night). *N. benthamiana* plants were transformed as previously described (Gantner *et al.*, 2019); a detailed protocol is provided online (dx.doi.org/10.17504/protocols.io.sbaeaie). Arabidopsis wild type accession Col was used, and plants were cultivated under short day conditions (8h light, 23/21°C day/night, 60 % relative humidity) or in a greenhouse under long day conditions (16h light) for seed set. Arabidopsis was transformed by floral dipping as previously described (Logemann *et al.*, 2006). Quartz sand plates for selection with 5-FC were prepared as previously described (Davis *et al.*, 2009).

### Molecular cloning and selection of tomato U6/U3 promoter fragments

The GoldenGate technique following the Modular Cloning syntax for hierarchical DNA assembly was used for most clonings (Engler *et al.*, 2008, Weber *et al.*, 2011). Previously reported plasmids belonging to the Modular Cloning Toolkit and the MoClo Plant Parts I and II collections were used (Engler *et al.*, 2014, Gantner *et al.*, 2018). For domestication of new DNA modules, respective fragments were amplified using Polymerase X (Roboklon), and ligated into Level 0 vectors.

*Sl*U6/U3 promoter fragments were selected from multiple sequence alignments of U6/U3 genes and upstream regions from Arabidopsis and tomato. Promoter fragments encompassing approximately 250 nt and 120 nt upstream of the predicted transcription start site were cloned, and functionally verified by expression of sgRNAs in reporter-based assays. No functional differences were observed, and shorter promoter fragments were used to generate data presented in this study. Shuttle vectors containing promoter fragments were cloned as previously described (Ordon *et al.*, 2017). Briefly, the promoter fragment and a second fragment encompassing a *ccdB* cassette and the sgRNA scaffold were PCR-amplified (*e.g.*, using oligonucleotides JS1744/1745 and STS24/JS1746 in case of the *Sl*U3 promoter), and subsequently fused by SOE-PCR. The SOE-PCR product was cloned into pUC57-BsaI in a cut/ligation reaction using *Eco*RV to yield the M1E module. This served as PCR template to amplify fragments for further shuttle vectors, which were (after *Dpn*I digestion) cloned as before. Plasmids and oligonucleotides are summarized in Table S1. Additional plasmids are listed in Appendix 1 as part of the cloning manual of our toolkit. Vector maps (GenBank) are provided in Appendix 2.

### Agroinfiltration, reporter-based nuclease activity assays and localization studies

For transient expression of proteins in *N. benthamiana* leaf tissues (agroinfiltration), respective T-DNA constructs were transformed into *Agrobacterium* strain GV3101 pMP90. Plate-grown bacteria were resuspended in Agrobacterium Infiltration Medium (AIM; 10 mM MES pH 5.8, 10 mM MgCl_2_), and infiltrated with a needleless syringe at an OD_600_ = 0.3 per strain. For qualitative determination of GUS activity, leaf discs were taken 3 dpi, stained with GUS staining solution (10 mM phosphate buffer pH 7, 10 mM EDTA, 1 mM potassium ferricyanate, 1 mM potassium ferrocyanate, 0,1 % Triton X-100, 0,1% X-Gluc) for 3 – 5h, destained with ethanol and dried in cellophane. Quantitative determination of GUS activity was done as previously described (Ordon *et al.*, 2017). Briefly, leaf material was ground in liquid nitrogen, powder resuspended in extraction buffer and cleared lysates prepared by centrifugation. Protein extracts were incubated with 4-methylumbelliferyl-beta-D-glucuronide (4MUG; 5 mM) for 1 – 2h, and MU production measured on a Tecan plate reader. Live-cell imaging was done using a Zeiss LSM780 confocal laser scanning microscope. GFP was excited using the 488 nm laser, and the detector range was set to 493-532 nm.

### Transgene counter-selection and genotyping

A motorized SteREO Discovery.V12 microscope (Zeiss) with UV illumination and an RFP filter set connected to an AxioCam MRc camera was used for selection of non-fluorescent seeds. Arabidopsis seeds containing the FAST marker (Shimada *et al.*, 2010, Engler *et al.*, 2014) were sorted by direct observation through the eyepiece. Weakly fluorescent *N. benthamiana* seeds containing the p2S3:mCherry/RFP cassette were imaged using the camera with an exposure time of approximately 5 s. Seeds were aligned on a wet sheet of Whatman paper, imaged and sorted. Selected seeds were directly sown into potting soil. *N. benthamiana* seeds containing *Sl*-FAST2 showed strong fluorescence, but fluorescence only developed when seeds were rehydrated, *e.g.*, by placing on wet Whatman paper. Non-fluorescent seeds containing *Sl*-FAST2 were selected by direct observation through the eyepiece. For selection via the *Ca-Bs3* marker, 3 – 4 weeks-old soil-grown segregants were infiltrated with a *Pseudomonas fluorescens* strain containing a chromosomal integration of the *Pseudomonas syringae* type III secretion system (“EtHAn”; Thomas *et al.*, 2009) and a plasmid for expression and translocation of the *Xanthomonas campestris* pv. *vesicatoria* transcription activator-like effector AvrBs3 (Gantner *et al.*, 2018). Plate-grown *P. fluorescens* bacteria were resuspended in 10 mM MgCl_2_, and infiltrated at an OD_600_ = 0.6. Cell death and development of the hypersensitive response was scored 2-3 dpi. DNA was extracted from Arabidopsis and *N. benthamiana* by the CTAB method, and Taq polymerase or Polymerase X (Roboklon) was used for genotyping.

### Multiplex editing and phenotypic analyses in Arabidopsis

Genome editing constructs were assembled as described in Appendix 1. sgRNA target sites were selected using ChopChop (for Arabidopsis; Labun *et al.*, 2016) or CRISPR-P (for N. benthamiana; Liu *et al.*, 2017). Target sites, oligonucleotides used for sgRNA construction and oligonucleotides used for genotyping are listed in Table S1.

For phenotypic analyses in the T_1_ generation, primary transformants were selected by resistance to BASTA. Presence of trichomes (editing of *GLABROUS1)* was scored by visual inspection in the vegetative phase, and editing of *ERECTA* upon bolting.

In the T_2_ generation, non-transgenic seeds were selected by absence of seed fluorescence. Plants were cultivated under short day conditions, and phenotypes associated with inactivation of genes targeted by genome editing were accessed as following: Presence/absence of trichomes was evaluated for editing of ***GLABROUS1*** at 2-3 weeks. The ***too many mouths*** and ***forked1*** phenotypes were identified in de-stained first leaves. Destaining involved treatment of leaves with a 3:1 mix of ethanol and acetic acid for 2 hours, overnight incubation in 95% ethanol and 1 hour in 5 % NaOH at 60°C (modified from Steynen and Schultz, 2003). Leaves were then mounted in 50% glycerol and phenotyped *via* DIC microscopy utilizing an inverted AxioObserver with an AxiocamMRm-DDE camera. ZenBlue software was used for capturing images and controlling the microscope. Three weeks-old seedlings were infected with *Hyaloperonospora arabidopsidis* isolates Cala2 and Emwa1 to analyze editing of the ***RPP2a*** (Sinapidou *et al.*, 2004) and ***RPP4*** (van der Biezen *et al.*, 2002) loci, respectively, and infection phenotypes were scored 6 dpi by Trypan Blue staining as previously described (Stuttmann *et al.*, 2011). To evaluate functionality of ***RPS2*** (Debener *et al.*, 1991) and ***RPS5*** (Warren *et al.*, 1998), approximately 5 weeks-old plants were syringe-infiltrated with *Pseudomonas syringae* strain DC3000 derivatives containing plasmids for the expression of AvrRpt2 or AvrPphB, respectively, at an OD_600_=0.05. Plants were left covered over-night, and development of the HR was scored 16-24 hpi. Results of phenotyping are presented in Table S2.

### Amplicon sequencing and data analysis

Oligonucleotides for PCR-amplification (Table S1) were designed using the NCBI primer designing tool. DNAs from pooled plant material or individual plants were tested for absence of the T-DNA using zCas9i-specific oligonucleotides. Amplicons encompassing RGN-targeted regions were pooled, purified, quantified using a NanoDrop and sequenced by Genewiz (Amplicon-EZ). Paired-end raw reads were adapter- and quality- trimmed using Trim Galore with default parameters (https://github.com/FelixKrueger/TrimGalore, v0.4.0). Trimmed reads were merged (-minhsp 10 - fastq_minmergelen 20) and dereplicated (--strand both) using USEARCH (Edgar, 2010; v11.0.667_i86linux32). Merged, dereplicated reads were mapped to the sequences of the 15 amplicons using BWA mem with default parameters (Li and Durbin, 2010). Mappings were analyzed with samtools view (Li *et al.*, 2009; v1.2, using htslib 1.2.1) allowing to extract statistics for individual amplicons.

## Accession Numbers

Accession numbers for Arabidopsis genes are provided in Table S1. Sequence information for *N. benthamiana* genes can be accessed on solgenomics.net.

## Acknowledgements

This work was supported by GRC grant STU 642-1/1 (Deutsche Forschungsgemeinschaft, DFG) to JS. UB is grateful for financial support by the Leibniz price from the DFG and the Alfried Krupp von Bohlen und Halbach Stiftung. FF was financed by an ERASMUS mobility program. We would like to thank Bianca Rosinsky for great work in the greenhouse. We are grateful to Jane Parker and Kenichi Tsuda for providing *Pseudomonas syringae* strains. Laurent Nussaume and Nathalie Leonhardt are acknowledged for providing seeds of 35S:FCY.UPP transgenic lines. Matthieu Joosten, Wen Huang and Sergio Landeo Villanueva selected genes for multiplex-editing in *N. benthamiana*.

## Supporting Information

**Supplemental Figure S1**: Functional verification of pDGE nuclease vectors and U6/U3 promoters.

**Supplemental Figure S2**: Efficiency of *Agrobacterium*-mediated transformation of *Nicotiana benthamiana* using pDGE vectors.

**Supplemental Figure S3**: Transgene counter-selection in *N. benthamiana* by seed fluorescence and inducible cell death.

**Supplemental Figure S4**: Toxicity of 5-FC to Arabidopsis plants on quartz sand plates.

**Supplemental Figure S5**: Toxicity of 5-FC to *N. benthamiana* on quartz sand plates.

**Supplemental Figure S6**: PCR-genotyping of transgene-free T_1_ *N. benthamiana* plants from multiplex-editing.

**Supplemental Figure S7**: Sequencing of alleles from *N. benthamiana* plant 747-1-4.

**Supplemental Figure S8**: Sequencing of alleles from *N. benthamiana* plant 747-1-6.

**Supplemental Figure S9**: Sequencing of alleles from *N. benthamiana* plant 747-3-8.

**Supplemental Figure S10**: Sequencing of alleles from *N. benthamiana* plant 747-3-9.

**Supplemental Figure S11**: Verification of T-DNA integrity in additional Arabidopsis T_1_ transformants.

**Supplemental Figure S12**: Details on alleles detected in individual Arabidopsis T_2_ segregants by amplicon sequencing.

Supplemental Table S1: Plasmids and oligonucleotides used in this study.

Supplemental Table S2: Phenotypic analysis of Arabidopsis T_2_ segregants from multiplex editing.

Appendix 1: Cloning manual for pDGE vector system.

Appendix 2: Annotated sequence files for pDGE vectors.

## Author Contributions

KB, PM, JO, JE, JG, RH and CK performed experiments, analyzed data and contributed to preparation of figures. TB and JK analyzed amplicon sequencing data. SM provided the intron-optimized Cas9 module prior to publication, and discussed data. JS designed the study, performed experiments, analyzed data, supervised experimental work, prepared final figures, and wrote the manuscript with contributions from JE and all authors.

---

**Supplemental Figure S1:**
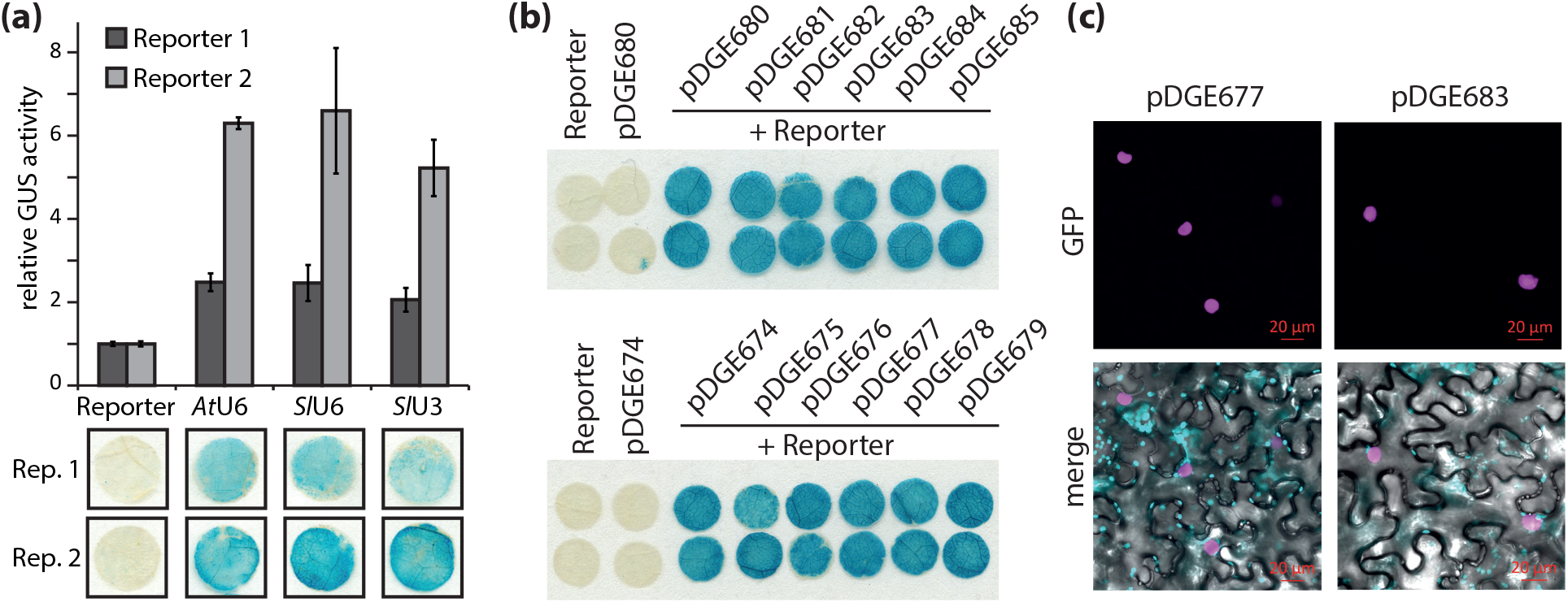
Functional verification of pDGE nuclease vectors and U6/U3 promoters. **(a)** Comparison of nuclease efficiency upon sgRNA expression by *At*U6, *Sl*U6 or *Sl*U3 promoter fragments using ß-Glucuronidase (*GUS*) nuclease activity reporters. Reporters consist of a 35S promoter-controlled *GUS* gene, which is shifted out-of-frame by an insertion subsequent to the initiating ATG (Ordon et al., 2019). Nucleases target this insertion sequence, and repair of double-strand breaks leads to reconstitution of a functional *GUS* gene. *Agrobacterium* strains containing reporters with two different target sites (Reporter 1: ATCCCGGAATTATCAGCACGA**GGG**; Reporter 2: TATGCTGCATGTAATCTGAAA**GGG**; PAM in bold) were infiltrated into *N. benthamiana* leaves, alone or in combination with strains for nuclease expression. Nuclease constructs contained sgRNAs targeting the respective reporter construct under control of the indicated U3/U6 promoter fragments. GUS activity resulting from reporters alone was arbitrarily set to 1, and error bars represent standard deviations from four replicates. Leaf discs from qualitative GUS staining are shown below the graph. Leaf material was harvested 3 dpi. **(b)** Verification of activity of nucleases encoded by the indicated constructs via *GUS*-based reporter assays. The sgRNA targeting Reporter 2 was cloned into the indicated recipient vectors, and Agrobacterium strains containing these constructs were used in reporter assays as in (a). **(c)** Detection and localization of GFP-Cas9. Tissue samples from the same experiment as shown in (b) were analyzed by confocal laser scanning microscopy. GFP - magenta; chlorophyll A - cyan.

**Supplemental Figure S2:**
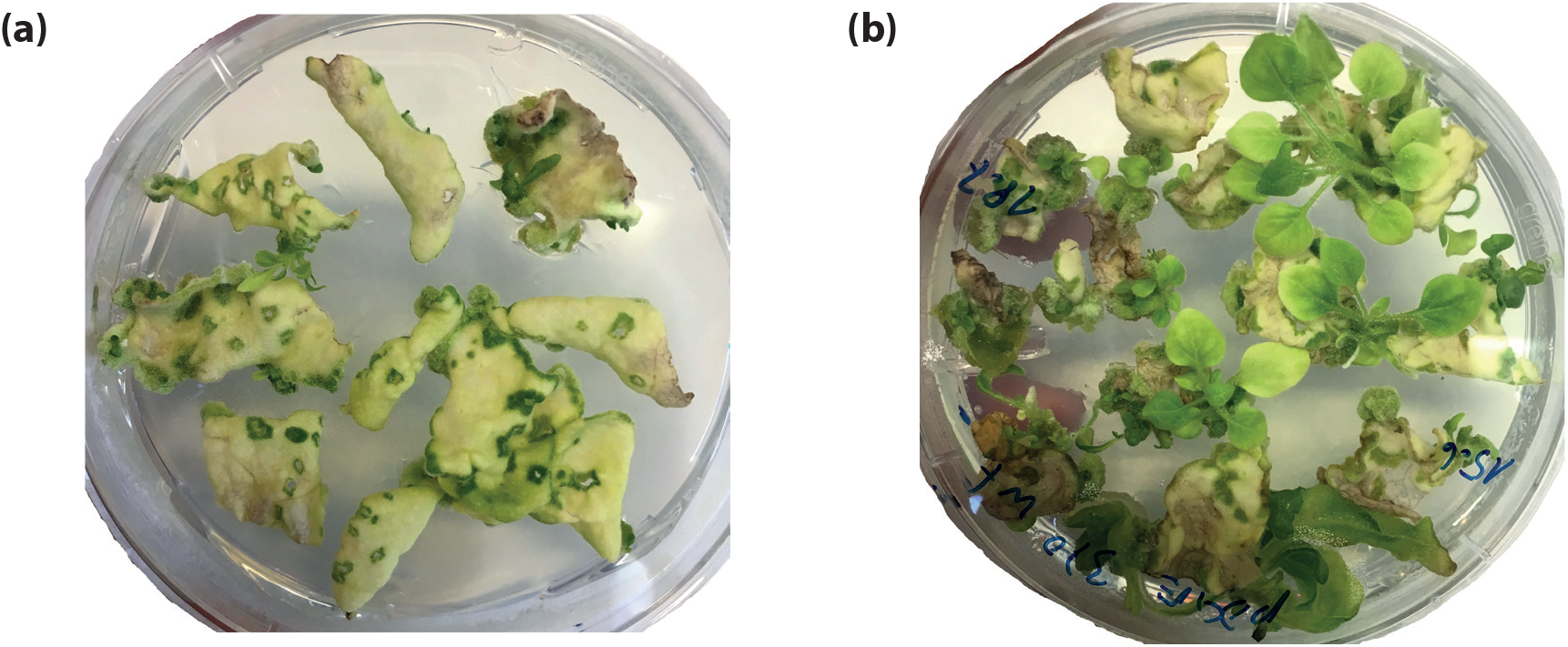
Efficiency of *Agrobacterium*-mediated transformation of *Nicotiana benthamiana* using pDGE vectors. Greenhouse-grown *Nicotiana benthamiana* plants were transformed as described in Materials and Methods. Four to five weeks after transformation (a), the formation of multiple calli from individual leaf sections became apparent. After 6-7 weeks (b), shoots developed from most calli.

**Supplemental Figure S3:**
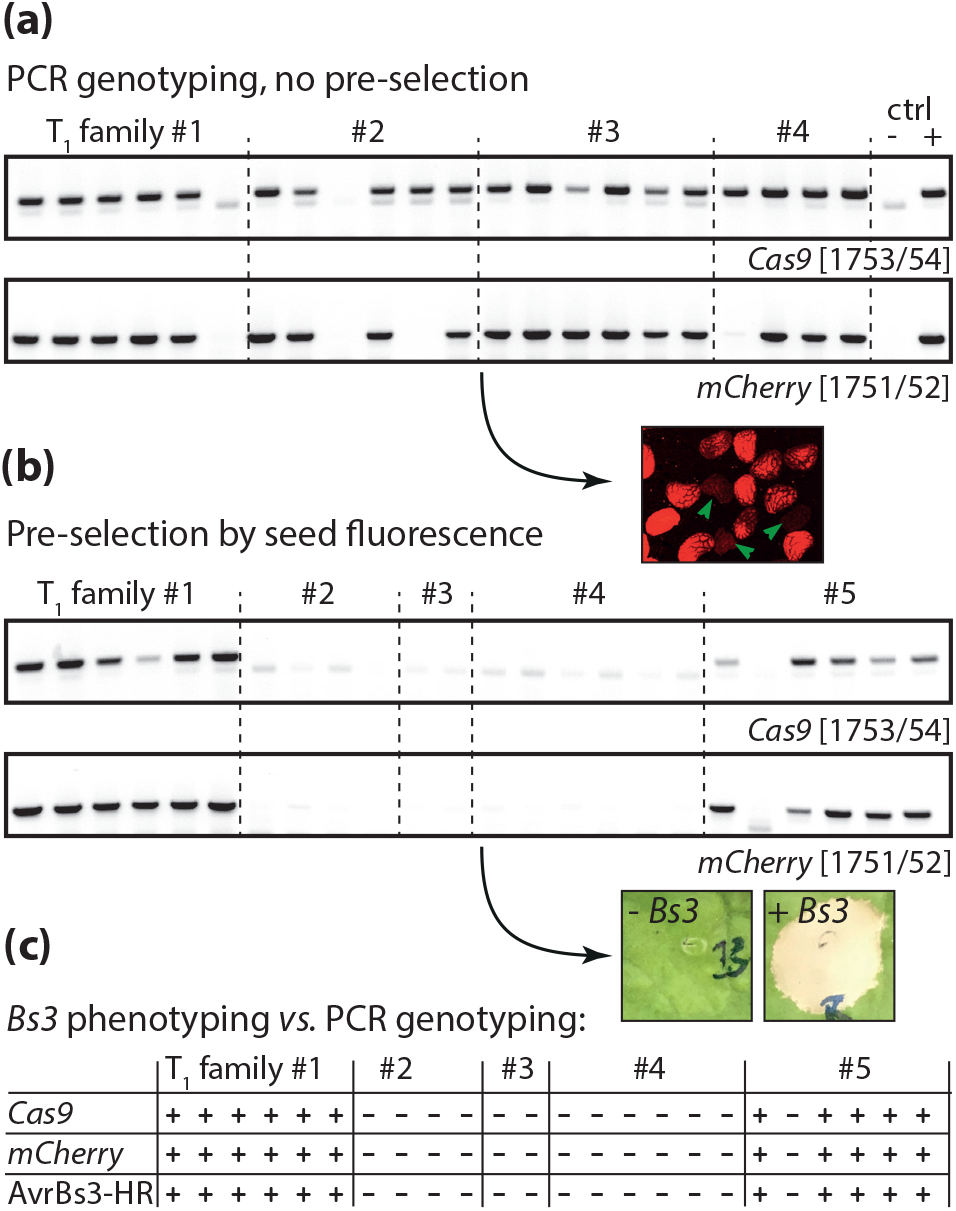
Transgene counter-selection in *N. benthamiana* by seed fluorescence and inducible cell death. **(a)** PCR-genotyping of randomly selected *N. benthamiana* T_1_ segregants with two different T-DNA-specific primer pairs. DNA of a wild type plant was used as negtive control (ctrl −), and DNA of a T_0_ regenerated plant served as positive control (+). **(b)** As in a), but seeds were pre-selected for absence of seed fluorescence. **(c)** Phenotyping of T_1_ segregants for presence of the *Bs3* marker gene. The same T_1_ segregants analyzed by PCR in b) were infiltrated with a *Pseudomonas fluorescens* derivative for translocation of AvrBs3. Appearance of cell death (see inset) was evaluated 3 dpi, and results were summarized in a table in comparison to results from PCR-genotyping.

**Supplemental Figure S4:**
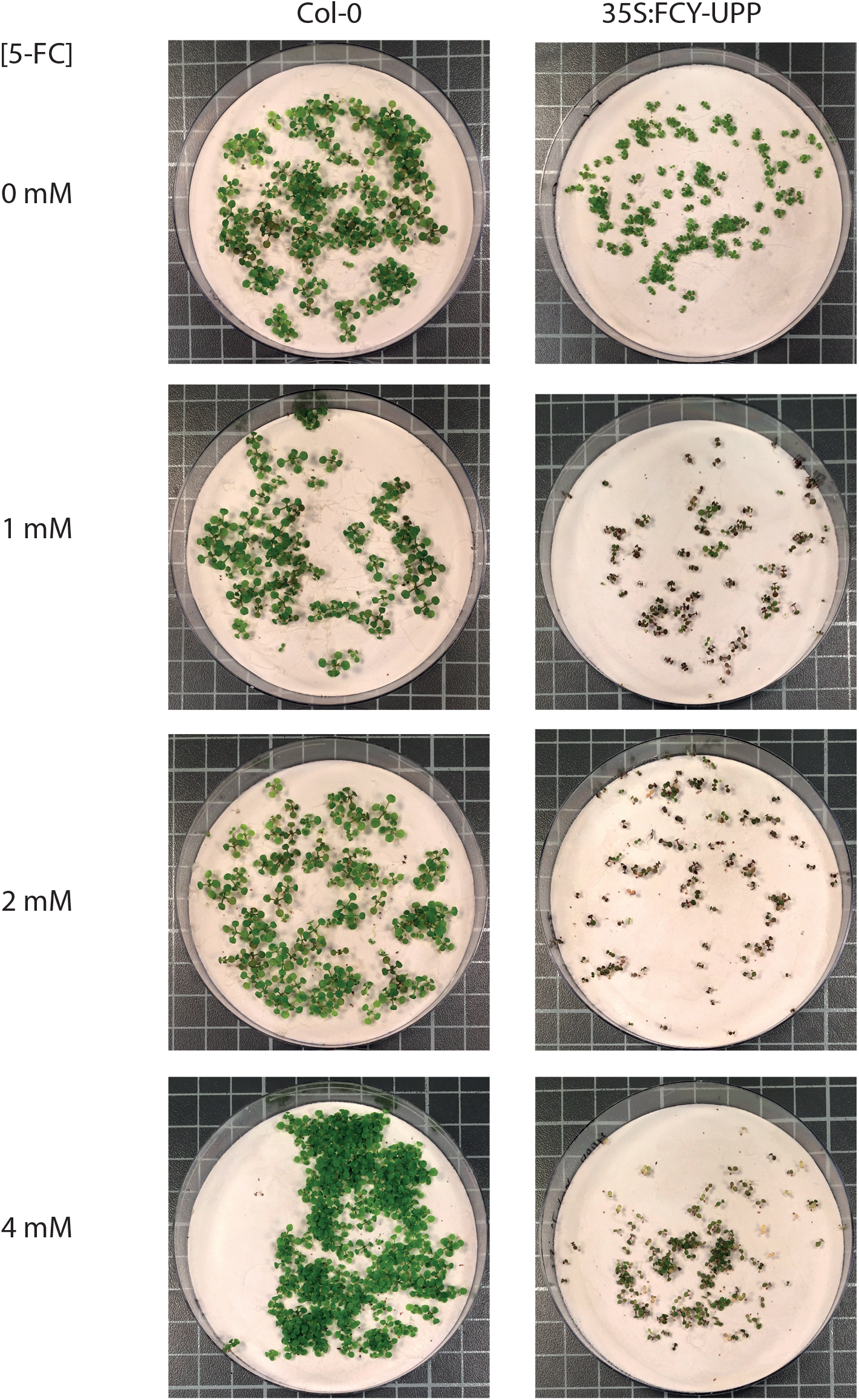
Toxicity of 5-FC to Arabidopsis plants on quartz sand plates. Arabidopsis wild-type or FCY-UPP transgenic plants (Leonhardt et al., 2020) were grown on quartz sand plates supplemented with 1/4 MS solution containing the indicated concentrations of 5FC. Plates were documented after 18 days.

**Supplemental Figure S5:**
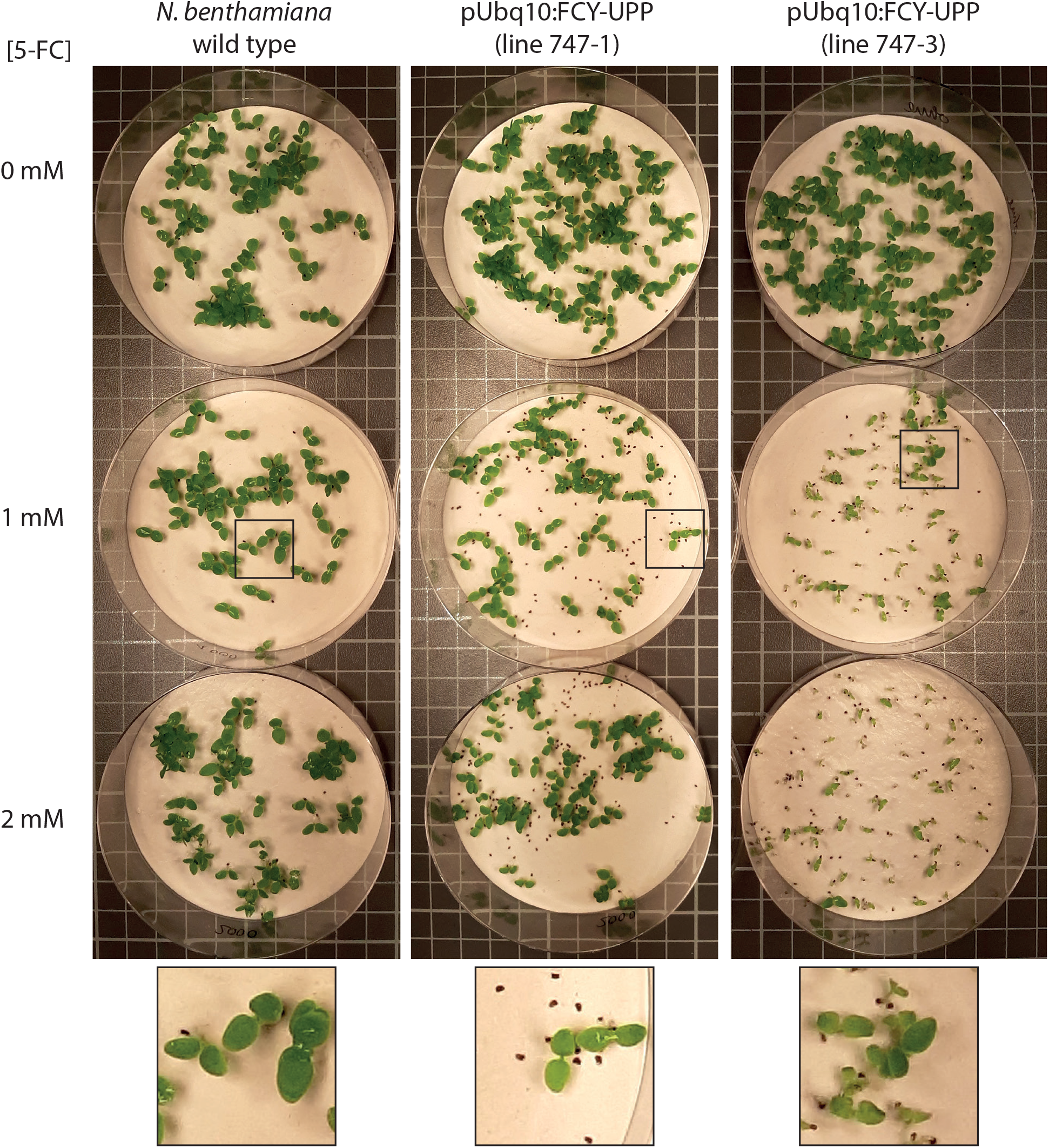
Toxicity of 5-FC to *N. benthamiana* on quartz sand plates. *N. benthamiana* wild-type or transgenic plants containing a derivative of pDGE792, an editing vector containing the pUbq10:FCY-UPP cassette as selection marker, were grown on quartz sand plates supplemented with 1/4 MS solution containing the indicated concentrations of 5-FC. Plates were documented after 12 days. A magnification of the area marked by a square of the 1 mM 5-FC plates is shown at the bottom.

**Supplemental Figure S6:**
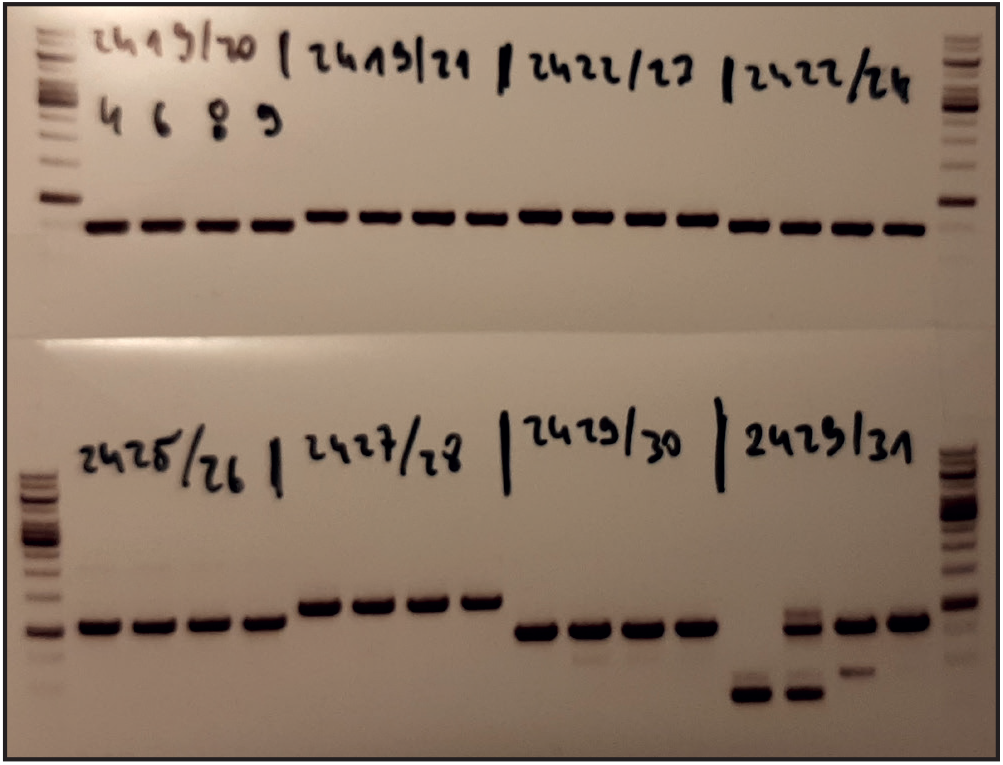
PCR-genotyping of transgene-free T_1_ *N. benthamiana* plants from multiplex- editing. DNA from plants 747-1/4, 747-1/6, 747-3/8 and 747-3/9 was used for PCR amplification of loci targeted for editing in *N. benthamiana* with indicated oligonucleotides. Following associations between oligonucleotides and *N. benthamiana* gene models [oligonucleotide sequences]: 2419/2420: Niben101Scf02460g01004 [CCAAAACCAAAGGCAAAGGAG / GATTAGCCAGGTAGATAGTGG] 2419/2421: Niben101Scf06739g05004 [CCAAAACCAAAGGCAAAGGAG / GACTGAAGTTCCAAAAGCC] 2422/2423: Niben101Scf01445g02008 [GTGATGAAGATTGCATGGG / GTTTACGAGTTATTGCACGAT] 2422/2424: Niben101Scf02763g03011 [GTGATGAAGATTGCATGGG / CACCAACTCCCAAAAGCC] 2425/2426: Niben101Scf03268g03002 [GTGGTCATTCAAGGCTAGC / TTTTCTTAAACTCCGTGCCG] 2427/2428: Niben101Scf00870g13015 [TCATCATCATAGTAGGCTAGCC / TAGTCCTTTTGCAGCACCTA] 2429/2430: Niben101Scf10055g01006 [CGTAGGTTGTCATTCTCGGA / TGTAACTTCTTGGGGTACGT] 2429/2431: Niben101Scf03619g00005 [CGTAGGTTGTCATTCTCGGA / TGCGTACCATCAAAACAACC]

**Supplemental Figure S7:**
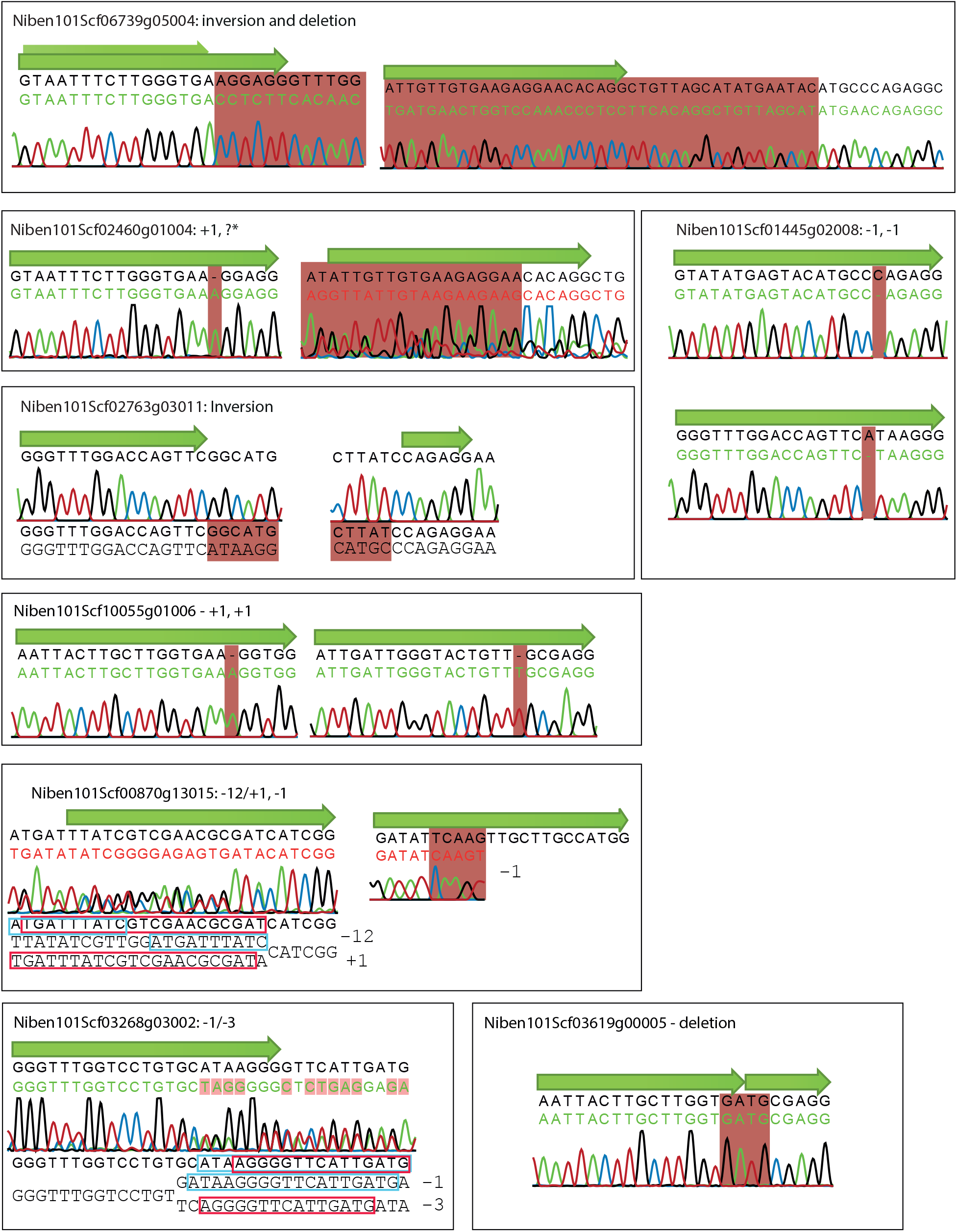
Sequencing of alleles from *N. benthamiana* plant 747-1-4. Sections of trace files from Sanger-sequencing of PCR products covering target sites in plant 747-1-4 are shown. Green arrows mark sgRNA target sites. In total, 29/30 analyzed target sites were most likely edited. Sequences were aligned using CLC Main Workbench (Qiagen). *analysis of the chromatogramm in later sections, where trace quality improved, suggested a heterozygous −3 mutation. Accordingly, this was scored as −3/wt for the display in Figure 3c.

**Supplemental Figure S8:**
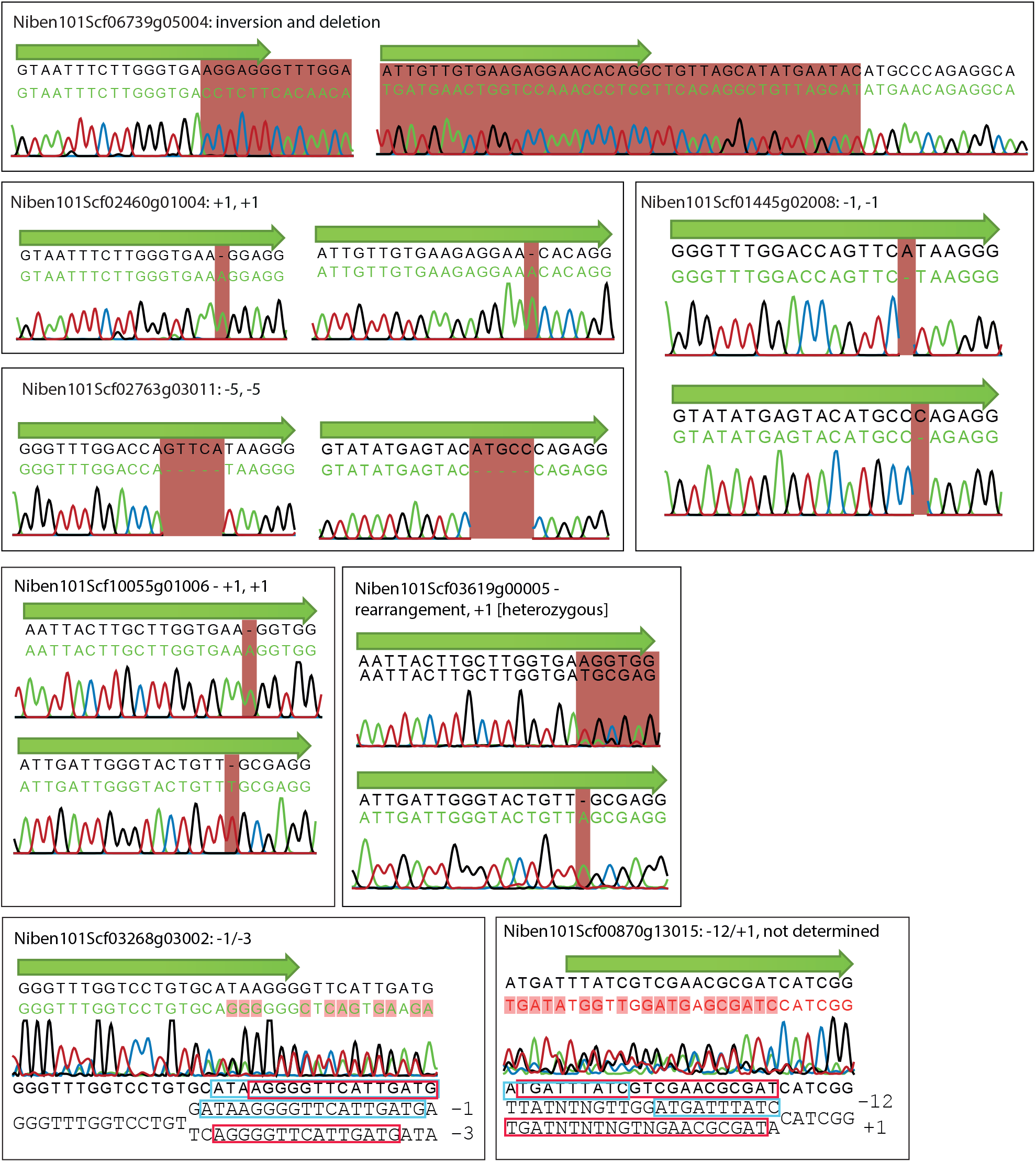
Sequencing of alleles from *N. benthamiana* plant 747-1-6. Sections of trace files from Sanger-sequencing of PCR products covering target sites in plant 747-1-6 are shown. Green arrows mark sgRNA target sites. In total, 28/28 analyzed target sites were edited. Sequences were aligned using CLC Main Workbench (Qiagen).

**Supplemental Figure S9:**
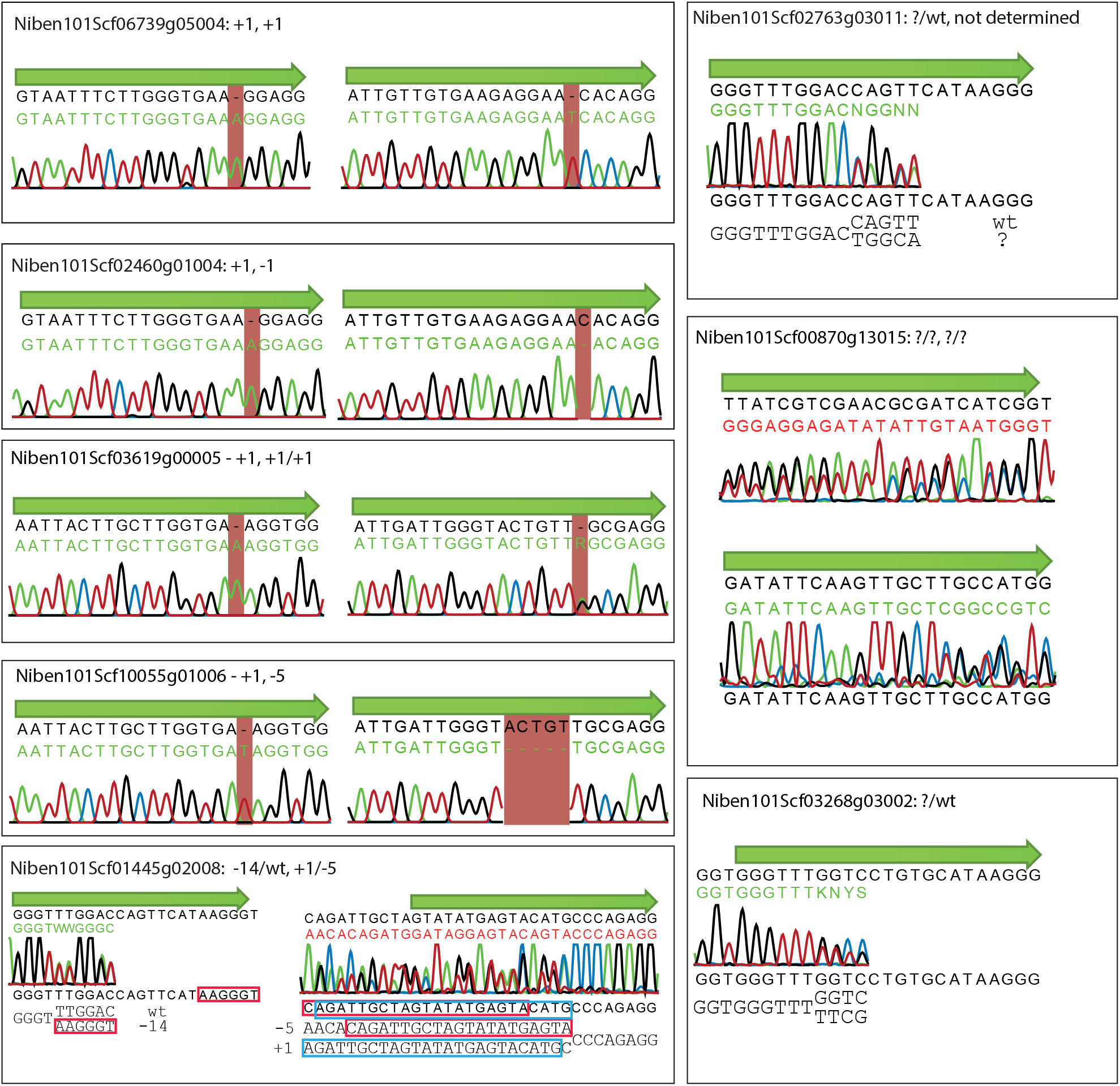
Sequencing of alleles from *N. benthamiana* plant 747-3-8. Sections of trace files from Sanger-sequencing of PCR products covering target sites in plant 747-3-8 are shown. Green arrows mark sgRNA target sites. In total, 25/28 analyzed target sites were edited. Sequences were aligned using CLC Main Workbench (Qiagen).

**Supplemental Figure S10:**
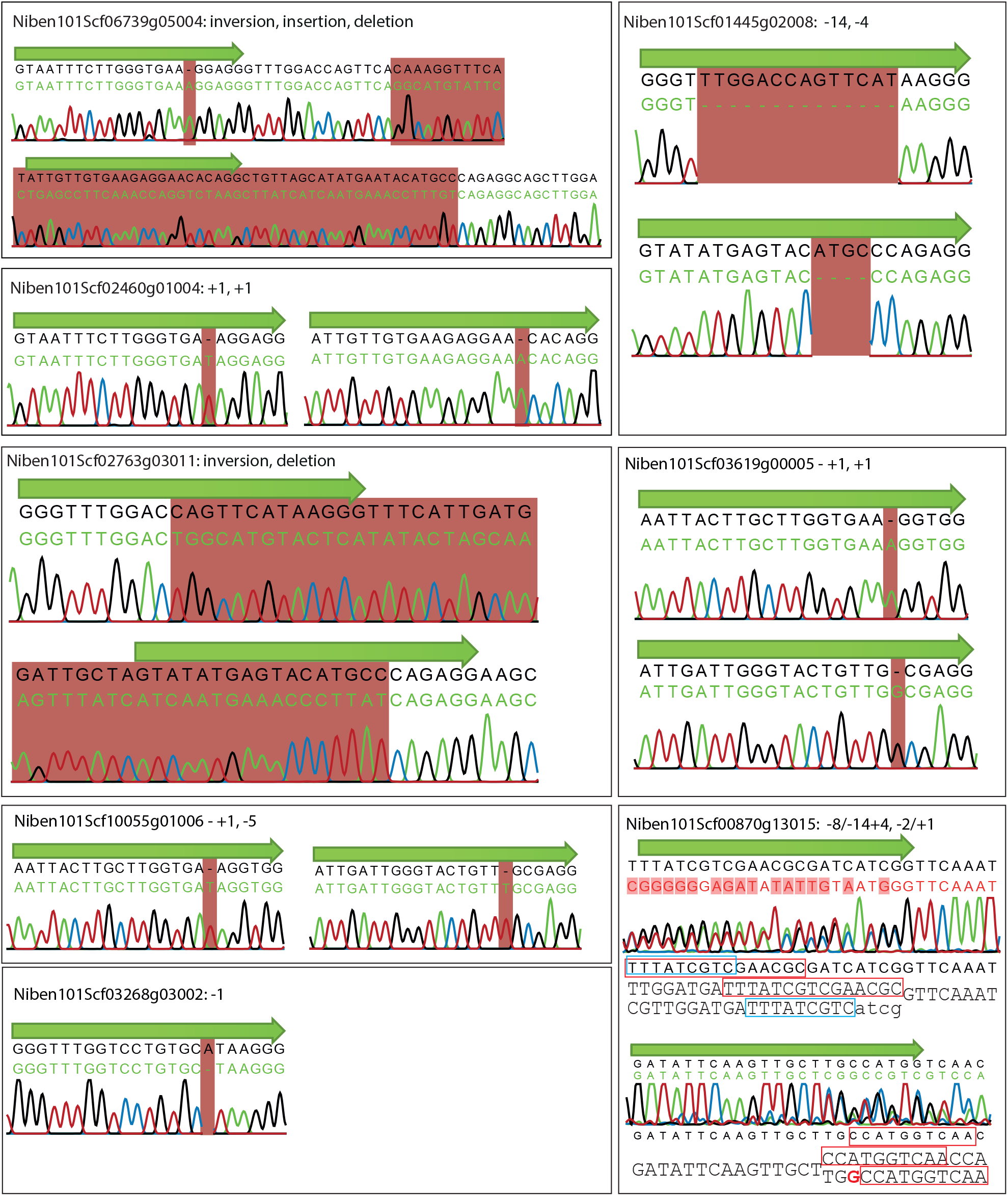
Sequencing of alleles from *N. benthamiana* plant 747-3-9. Sections of trace files from Sanger-sequencing of PCR products covering target sites in plant 747-3-9 are shown. Green arrows mark sgRNA target sites. In total, 30/30 analyzed target sites were edited. Sequences were aligned using CLC Main Workbench (Qiagen).

**Supplemental Figure S11:**
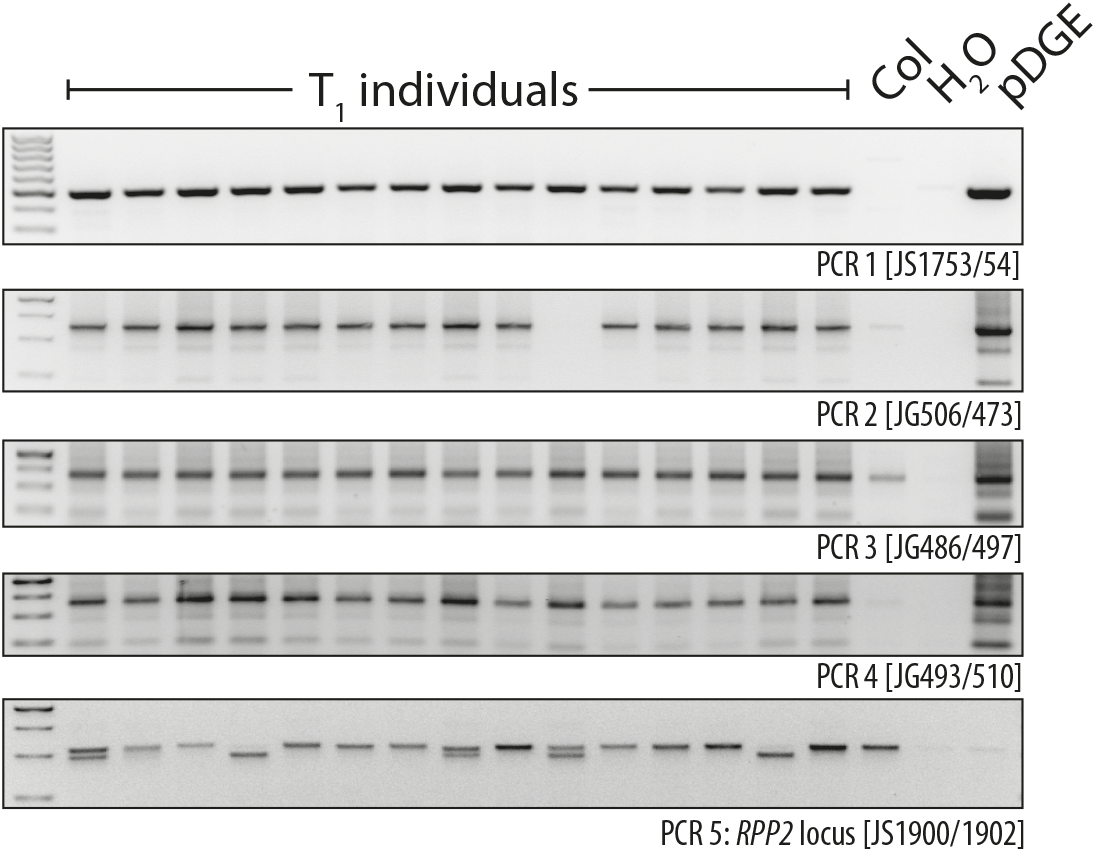
Verification of T-DNA integrity in additional Arabidopsis T_1_ transformants. As in Figure 4b, but additional transformants were tested. PCR1 - Cas9; PCRs 2/3/4 - different amplicons within the array of 24 sgRNA TUs; PCR5 - *RPP2* locus, also targeted by sgRNAs.

**Supplemental Figure S12:**
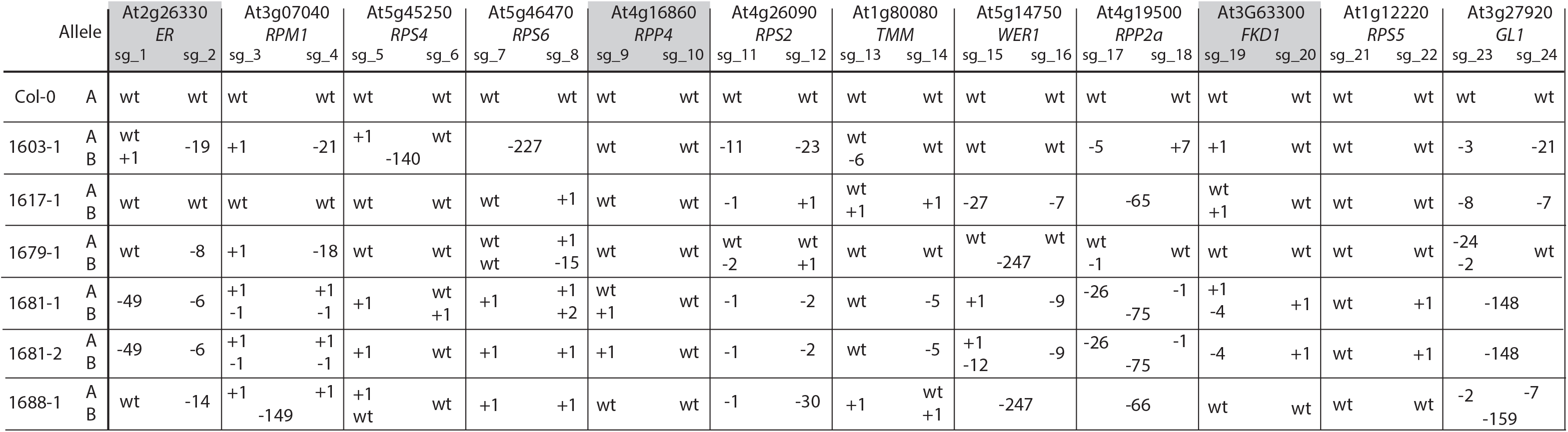
Details on alleles detected in individual Arabidopsis T_2_ segregants by amplicon sequencing. Target loci from indicated transgene-free T_2_ plants were PCR-amplified and analyzed by amplicon sequencing. Reads were quality filtered, merged and aligned to the Col reference genome to identify respective alleles. In most cases, analyzed PCR amplicons contained both target sites, thus allowing unambiguous assignment of alleles A and B. Target sites for loci marked by grey shading were PCR-amplified as two independent amplicons.

## pDGE Dicot Genome Editing vectors: Cloning Manual v5

### Shuttle vectors (Amp/Carb, Cm)

**sgRNA shuttle vectors with *Arabidopsis thaliana* U6 promoter: guide sequence overhangs ATTG-GTTT**

**sgRNA shuttle vectors with *Solanum lycopersicum* U6 promoter: guide sequence overhangs ATTG-GTTT**

**sgRNA shuttle vectors with *Solanum lycopersicum* U3 promoter: guide sequence overhangs GTCA-GTTT**

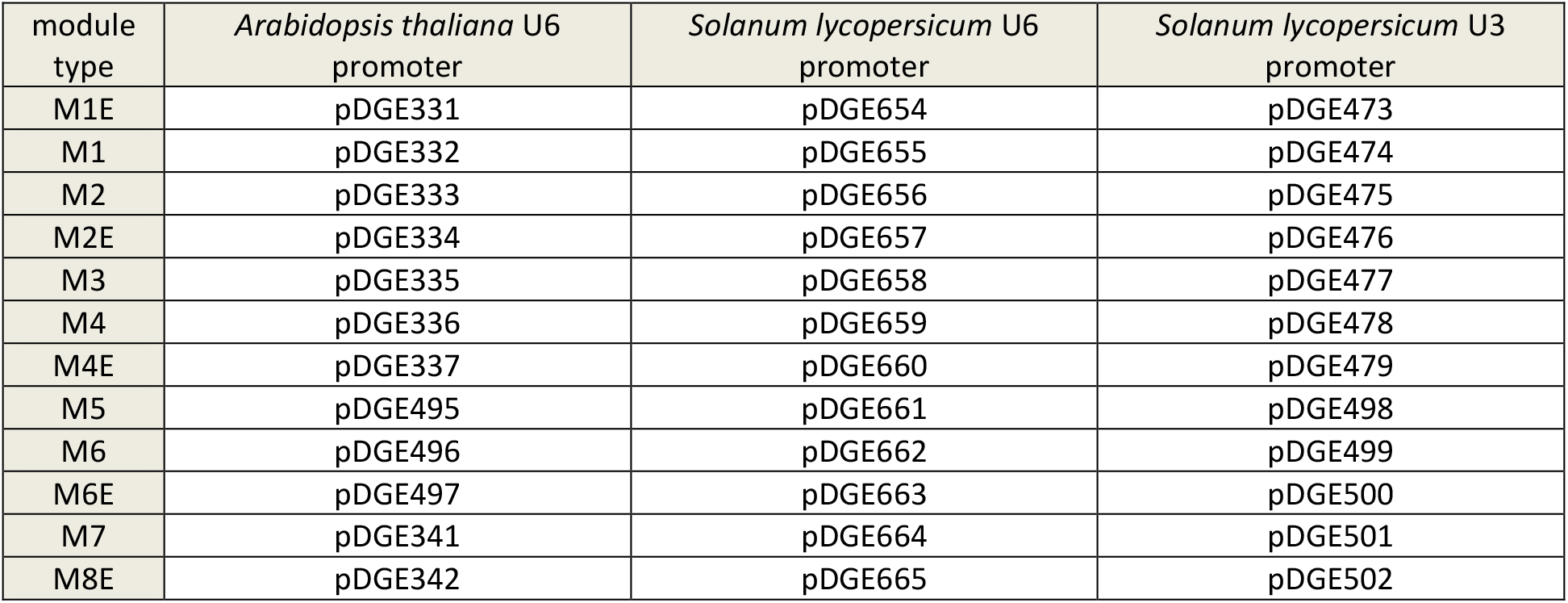

### Recipient vectors for *Arabidopsis thaliana* (pRPS5a:Cas9, FAST)

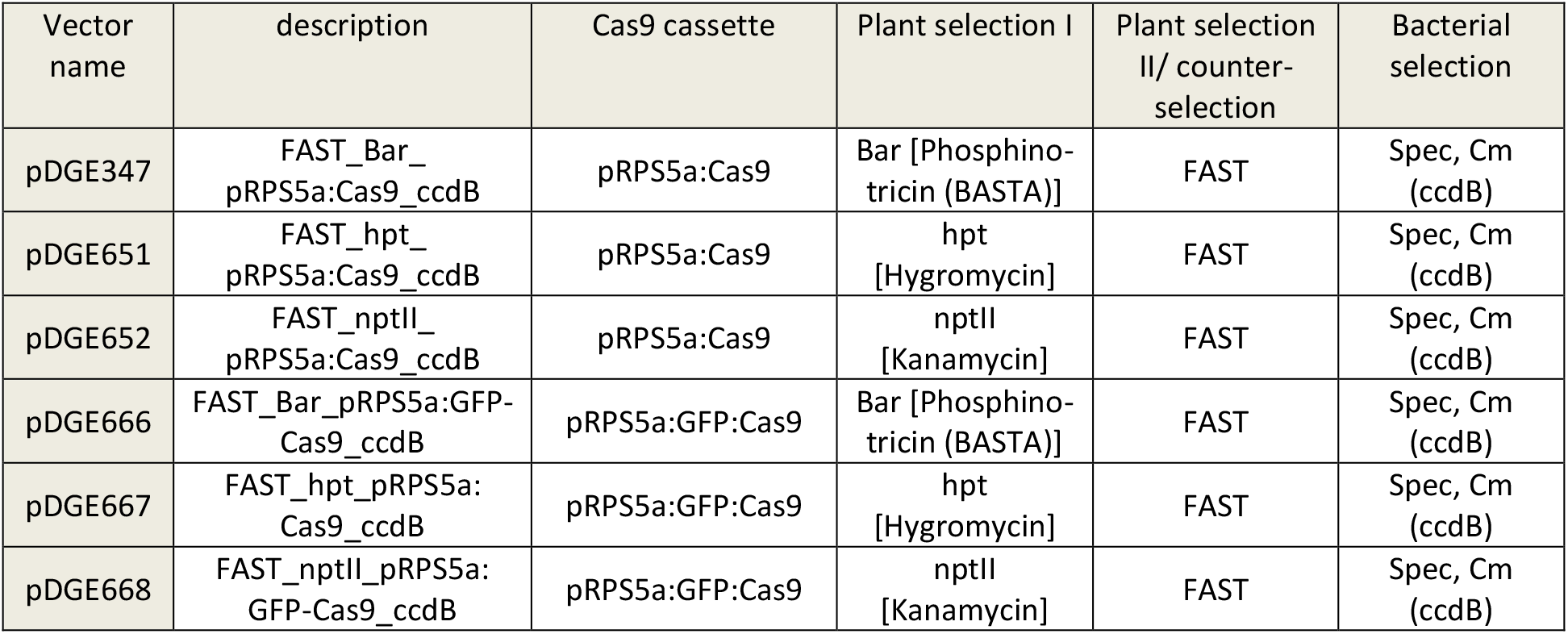

### Recipient vectors for *Nicotiana benthamiana* (p35S:Cas9 or pRPS5a:Cas9, secondary markers)

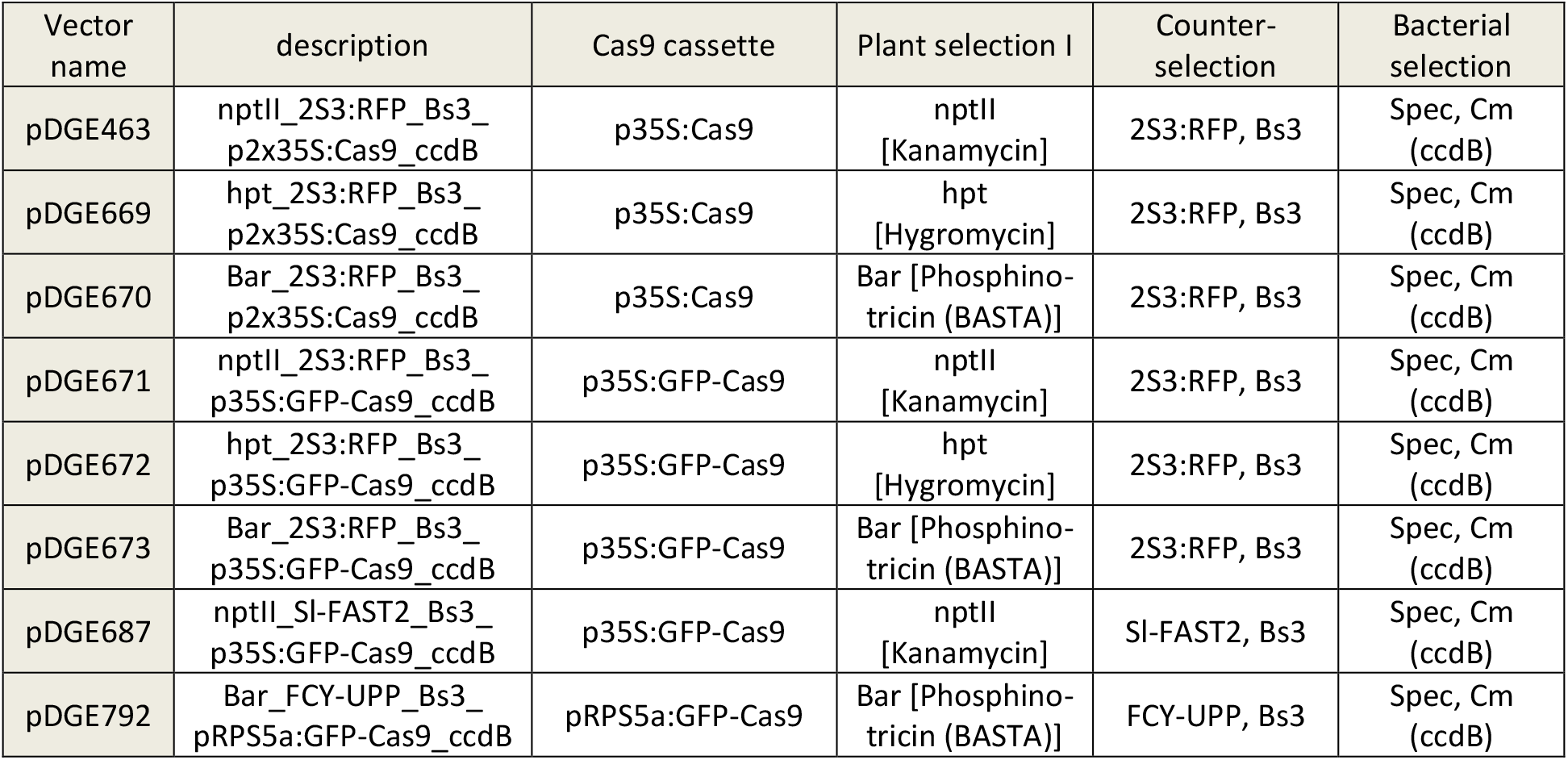

### Recipient vectors and end-linkers for higher order multiplexing

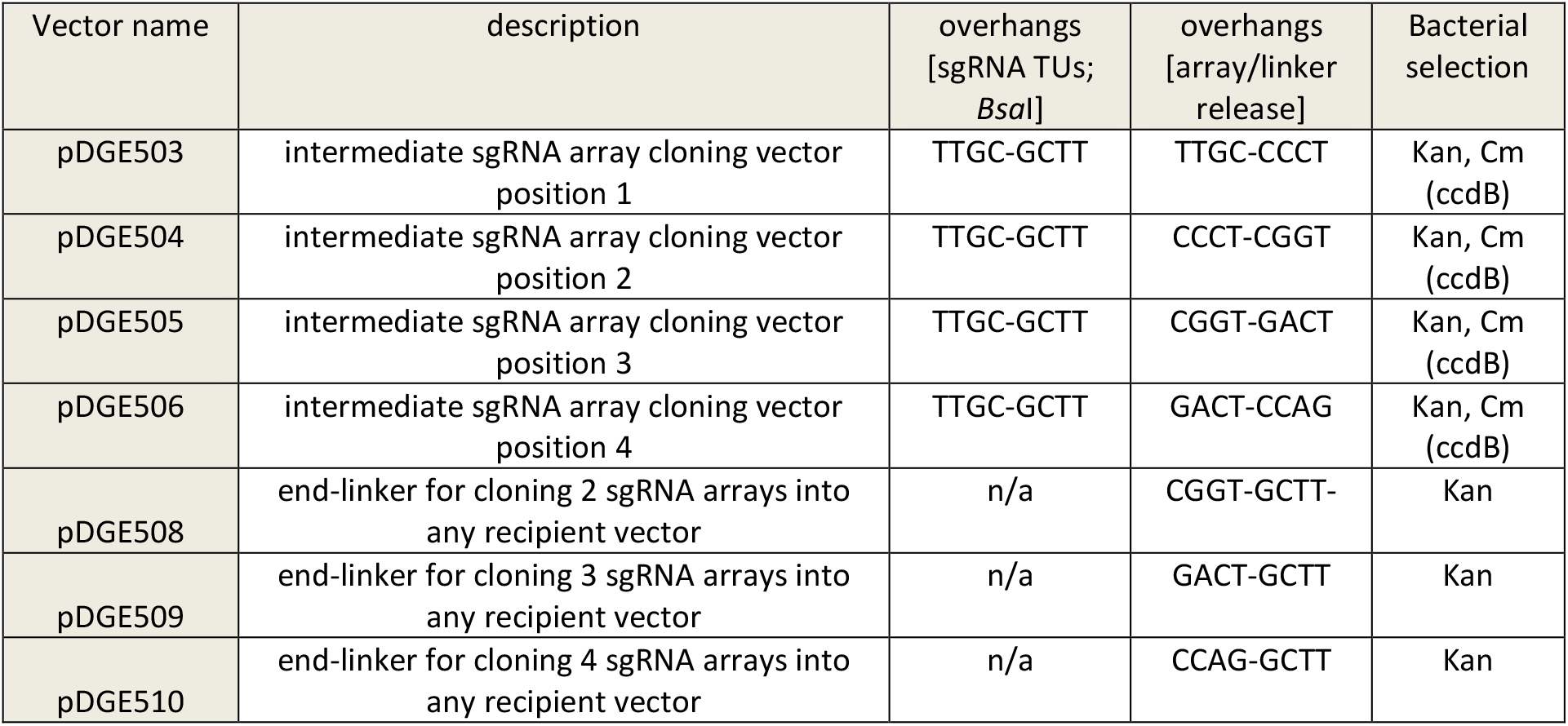

- Most empty vectors contain a *ccdB* cassette and must be propagated in DB3.1 or *ccdB* survival cells (Invitrogen / Thermo Scientific).
- Amp – Ampicillin, Carb – Carbenicillin, Spec – Spectinomycin, Kan – Kanamycin, Cm - Chloramphenicol
- Annotated sequence files are provided as a multi-record genbank file, which can be opened with any DNA sequence analysis software (e.g. CLC, Lasergene, Sequencher, Geneious, Vector NTI).

### Overall vector architecture

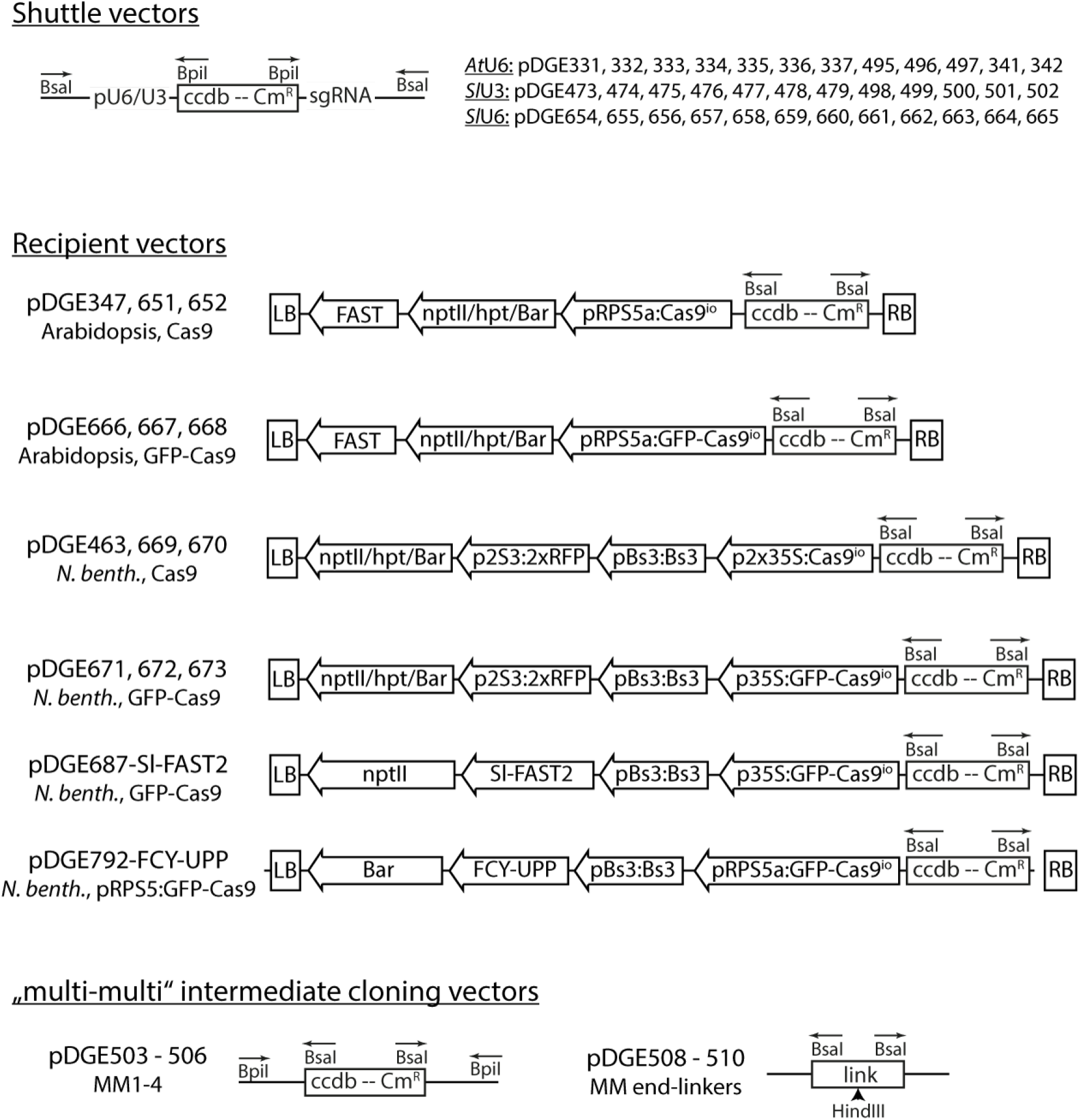

<u>**Note:**</u> The *Bsa*I fragment (prom-ccdB-sgRNA) of an M1E shuttle vector (pDGE331 (*At*U6), pDGE473 (*Sl*U3) or pDGE654 (*Sl*U6)) can be transferred into any recipient vector *via* a *Bsa*I GoldenGate reaction to obtain a novel recipient for rapid generation of single nuclease constructs. Such “one step, one nuclease” vectors (Ordon et al., 2017, Plant Journal) serve for direct cloning of hybridized oligonucleotides in the recipient, but do not offer multiplexing options. In the *Bsa*I GoldenGate reaction, both the empty recipient (*e.g.* pDGE347) and the assembly product (*e.g.* pDGE347 + *Bsa*I fragment from pDGE331) have the *ccdB* and *cat* (Chloramphenicol^R^) genes. It is therefore critical to add the M1E module in excess to the GoldenGate reaction (e.g. 10 fmol recipient, 40 fmol M1E), and to re-digest the GoldenGate reaction with *Bsa*I after a denaturation cycle, prior to transformation into *ccdB* survival cells.

## Selection and design of guide RNAs

sgRNA variable sequences are loaded into sgRNA shuttle vectors as 23-24 nt long, hybridized oligos in a *Bpi*I GoldenGate reaction. Phosphorylation of oligonucleotides is not required. Any online tool (such as *e.g.* <u>CHOPCHOP</u> or <u>CRISPR-P</u>) may be used for selection of sgRNAs, and cloning overhangs can be added manually or directly via the sgRNA design programs. Note that cloning overhangs differ for U3 and U6 promoter systems. Examples of target sites on plus and minus strands, and design of respective oligos to produce sgRNAs with the pDGE vector system, are provided below:

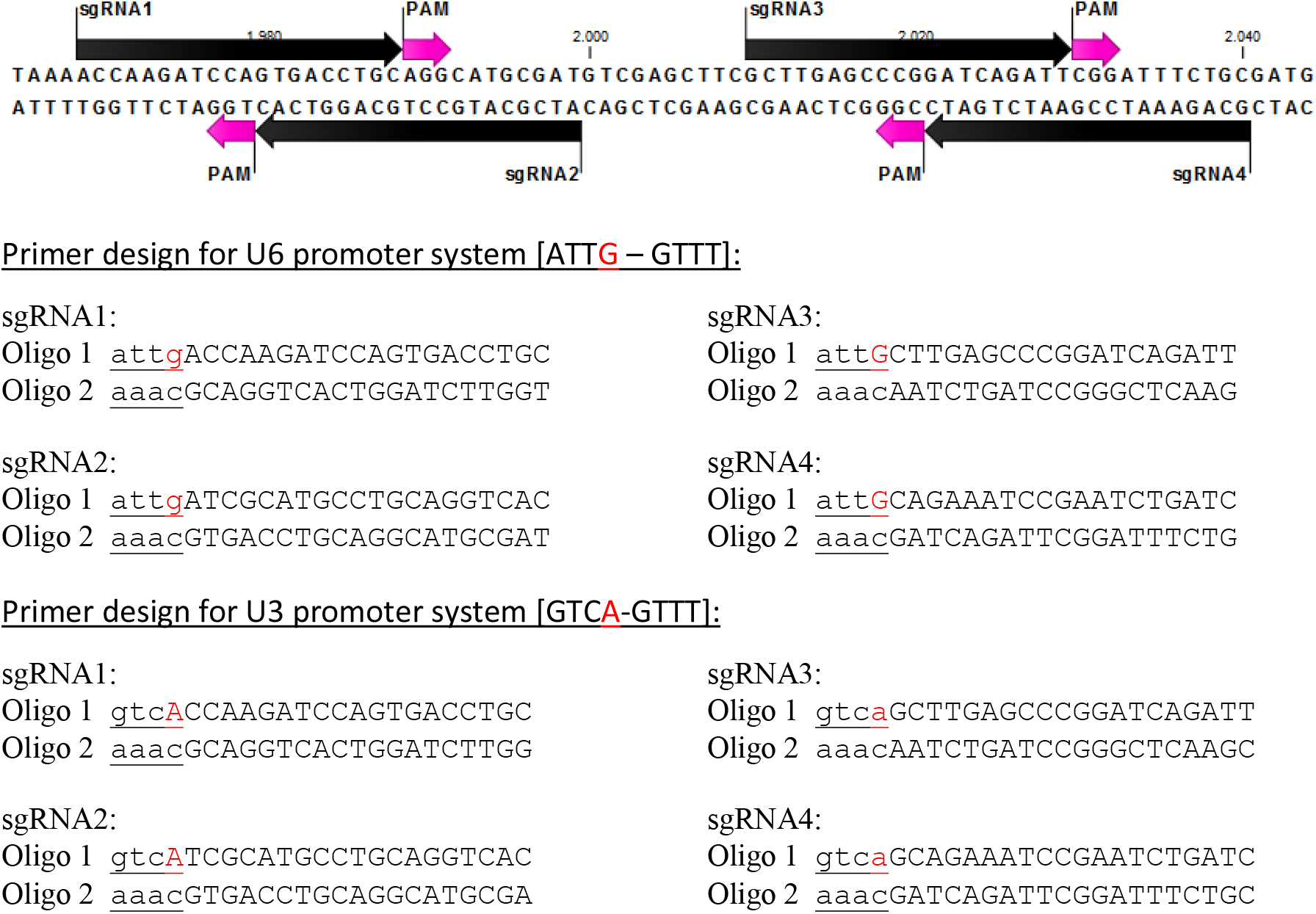

- The last nucleotide of the forward cloning overhang (marked in red) is the assumed transcription start site
- U6 promoter system: **sgRNAs 1/2** will contain 20 bases complementary to the target strand and a single non-complementary nucleotide in position 1. Oligonucleotides are 24 nt in length. **sgRNAs 3/4** will contain 20 bases complementary to the target strand. Oligonucleotides are 23 nt in length.
- U3 promoter system: **sgRNAs 1/2** will contain 20 bases complementary to the target strand. Oligonucleotides are 23 nt in length. **sgRNAs 3/4** will contain 20 bases complementary to the target strand and a single non-complementary nucleotide in position 1. Oligonucleotides are 24 nt in length.
- Avoid *Bsa*I sites [GGTCTC], *Bpi*I sites [GAAGAC] and polyT stretches [≥ 4 Ts; transcriptional termination] in sgRNA sequences.

### Cloning of sgRNA TUs in shuttle vectors

1. <u>Hybridization of oligonucleotides</u>

- oligonucleotide stock concentration: 100 μM
- mix oligos at 10 μM (for example 5 μl of each oligo + 40 μl H_2_O)
- denature oligos by heating to 98 °C for 5 min
- let cool down slowly (leaving tube @ RT for several minutes is sufficient)
- prepare a 1:200 dilution (50 fmol/μl) of the hybridized oligos
2. <u>Loading of oligonucleotides into sgRNA shuttle vectors</u>

cut/ligation reaction:

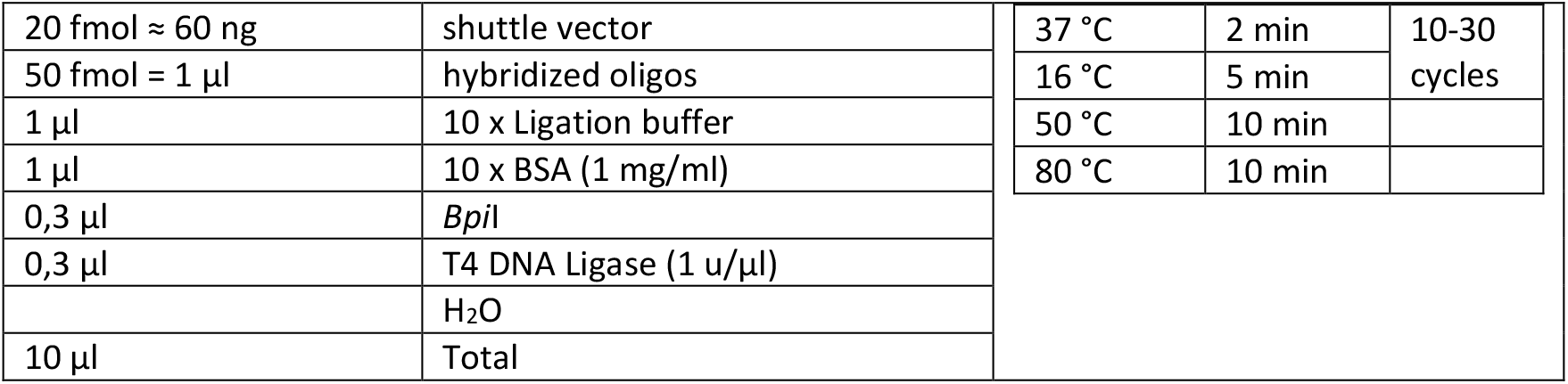

- transform cut/ligation reaction into Dh10b/TopTen cells
- optional: plate a 50 μl aliquot of the transformation on an LB-Carb plate
- directly inoculate liquid cultures (3 ml) with the transformed cells in selective media (LB-Carb)
- use liquid cultures for polyclonal plasmid preparation
- optional: perform test digest of the polyclonal plasmid preparations, *e.g. Pvu*II

## Assembly of plant transformation vectors containing up to eight sgRNA TUs

Derivatives of suitable shuttle vectors, loaded with sgRNA sequences, are used for assembly of one, two, four, six or eight sgRNA TUs in any recipient vector by a *Bsa*I cut/ligation reaction. This yields final constructs for plant transformation. Shuttle vectors from the different promoter systems are compatible. The following modules can be assembled:

1 sgRNA TU: M1E module

2 sgRNA TUs: M1 and M2E module

4 sgRNA TUs: M1, M2, M3 and M4E module

6 sgRNA TUs: M1, M2, M3, M4, M5, M6E module

8 sgRNA TUs: M1, M2, M3, M4, M5, M6, M7, M8E module

cut/ligation reaction:

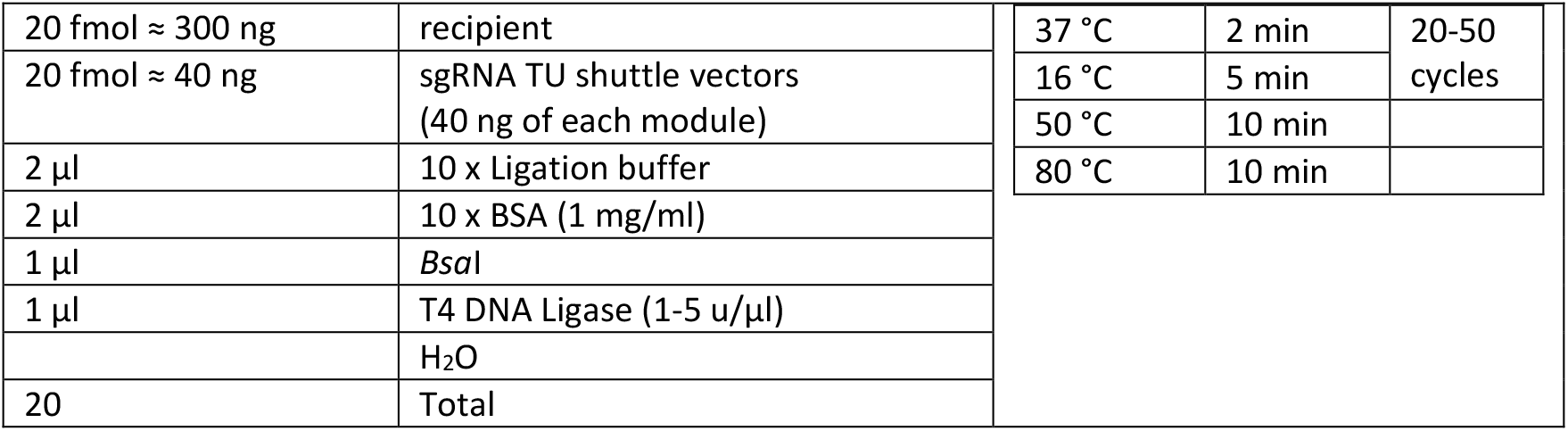

- transform cut/ligation reaction into Dh10b/TopTen cells
- plate on LB-Spec media
- 1-2 sgRNA TUs: Start liquid cultures from 2 clones
- 4-6 sgRNA TUs: Start liquid cultures from 2-3 clones
- 8 sgRNA TUs: Start liquid cultures from 3-4 colonies
- isolate plasmid DNA and verify by restriction digest
- confirm sgRNAarray by DNA sequencing: 1-4 sgRNA TU: oligonucleotide JS1132 6-8 sgRNA TUs: oligonucleotides JS1132 and M13f or JS2045 (when using pRPS5a:Cas9)
- transform plasmid into your favorite *Agrobacterium* strain

<u>useful oligonucleotides:</u>

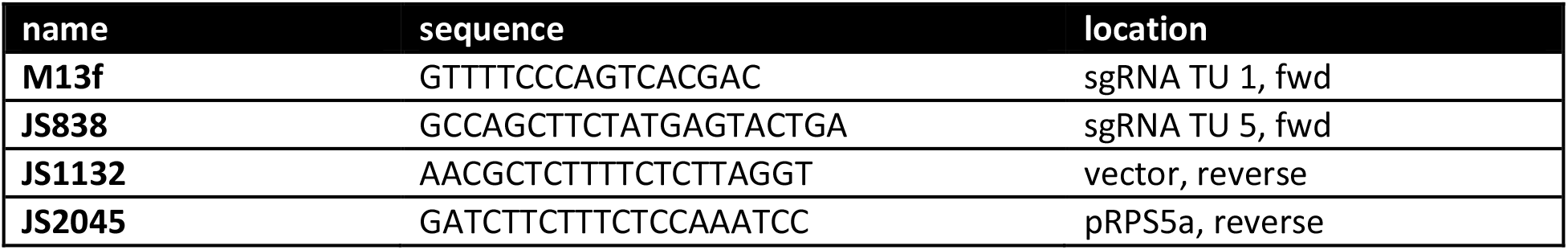

## Assembly of plant transformation vectors containing up to 32 sgRNA TUs

- sgRNA TUs, mobilized from shuttle vectors, are first assembled in intermediate “multi-multi” vectors as described above (section “Assembly of plant transformation vectors containing up to eight sgRNA TUs”). Multi-multi modules carry Kanamycin as selectable marker!

cut/ligation reaction:

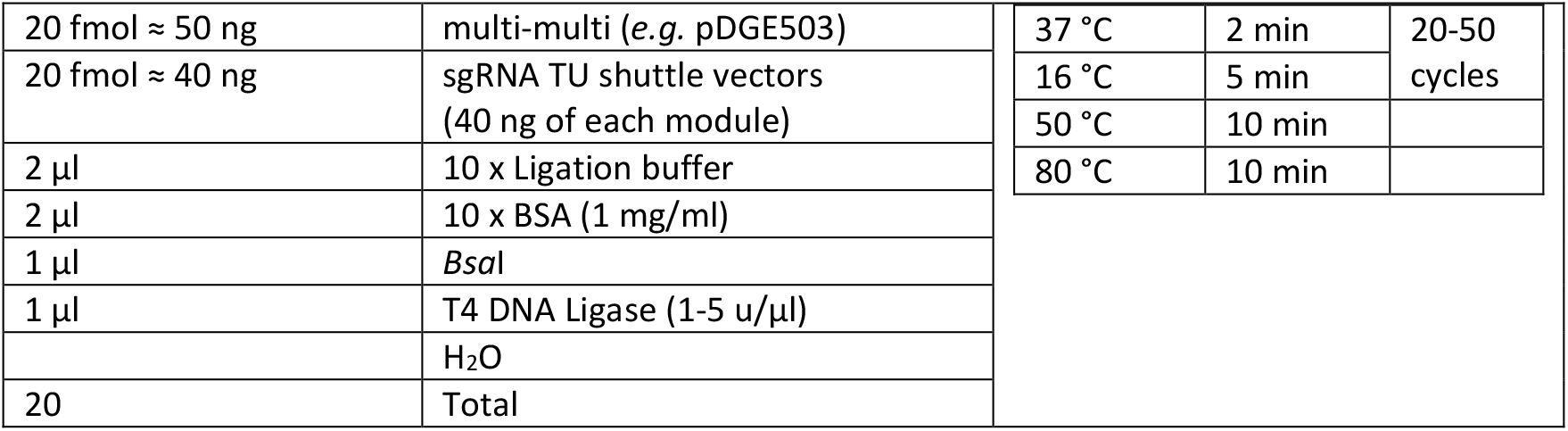

- Plasmid isolation, verification by restriction digest (*e.g. Bpi*I)
- sgRNAs arrays are verified by sequencing at this stage [oligonucleotides M13f or JS1691 and JS1692]
- sgRNA arrays are subsequently mobilized from multi-multi modules into any recipient vector in a GoldenGate reaction using both *Bsa*I and *Bpi*I (see next page).
- End-linkers are used to link the terminal overhang of a multi-multi module to the recipient vector.

The following assemblies are compatible:

up to 16 sgRNAs: derivatives of pDGE503 and pDGE504 together with pDGE508 (end-linker)
up to 24 sgRNAs: derivatives of pDGE503, 504 and 505 together with pDGE509 (end-linker)
up to 32 sgRNAs: derivatives of pDGE503, 504, 505 and 506 together with pDGE510 (end-linker)

cut/ligation reaction:

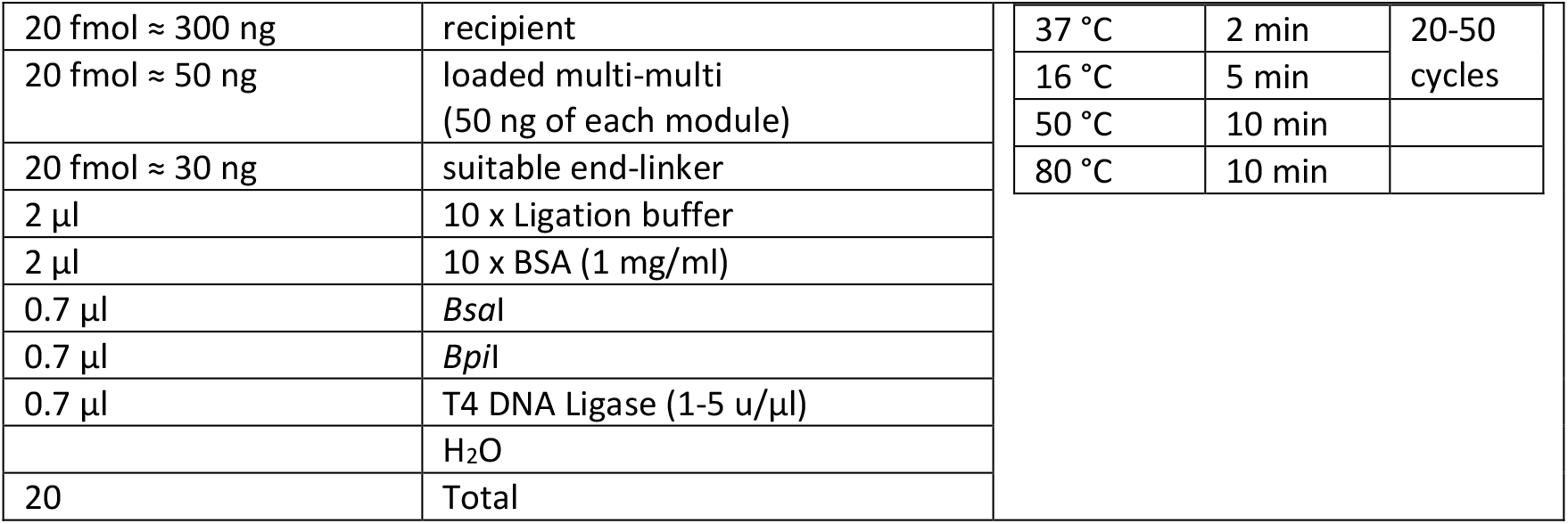

- transform cut/ligation reaction into Dh10b/TopTen cells
- plate on LB-Spec media
- start liquid cultures from 3-4 colonies
- isolate plasmid DNA and verify by restriction digest (e.g. *Hind*III)
- transform plasmid into your favorite *Agrobacterium* strain

<u>useful oligonucleotides:</u>

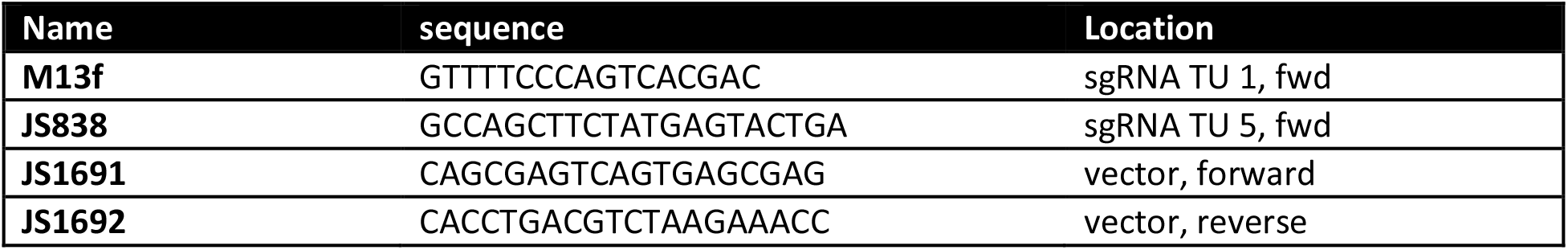

## Counter-selection using 5-fluorocytosine on quartz sand plates

**Material:**

- Quartz Sand [SIGMA 83340]
- MS salts including vitamins [Duchefa M0222]
- 0.7 M MES buffer, pH 5.7 [2-(N-morpholino)ethanesulfonic acid; adjust pH with KOH, sterilize by filtration]
- 5-fluorocytosine [SIGMA F7129; a stock concentration of 100 mM is possible, but requires careful redissolving upon thawing of frozen aliquots]

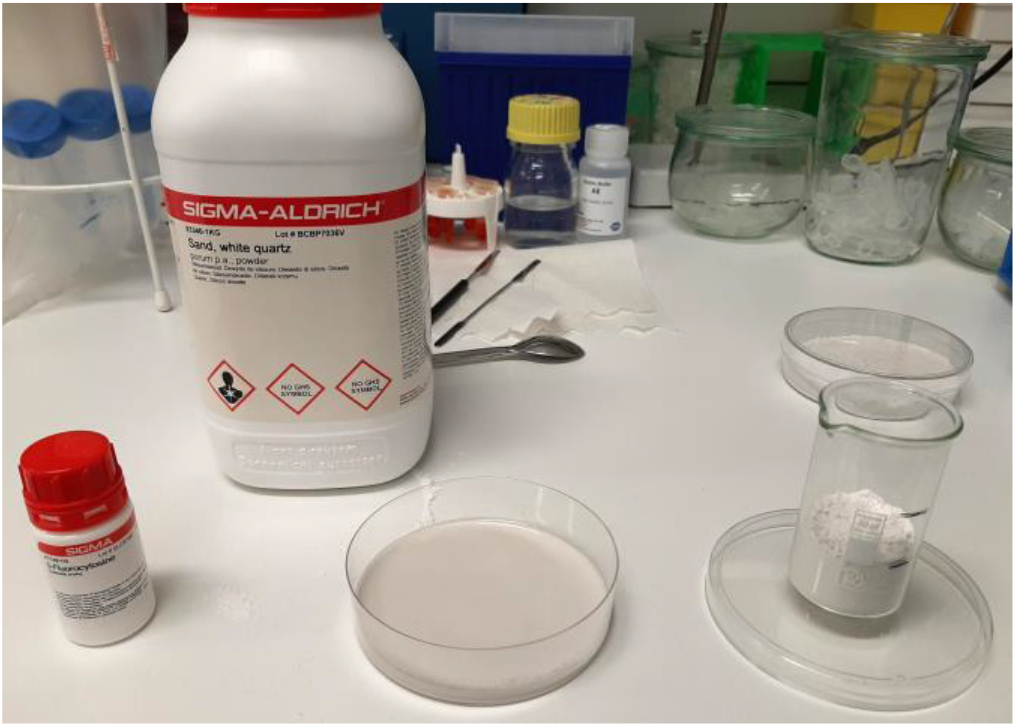

To prepare ¼ MS liquid medium: 20 ml MES buffer, 1.1 g MS salts on 1 l. Selective reagents should be added fresh, upon use of the media.

Cultivation and selection on sand plates was described previously (Davis *et al.*, 2009). Briefly, measure approximately 20 ml of sand, *e.g.* using a small beaker. Pour the sand into a petri dish and spread it using a spoon or spatula. Add 10 ml of media (1/4 MS containing 5-FC), mix by swiveling the plate to entirely imbibe the sand. Leave plate for a few minutes, and pour-off excess media. Sow seeds. During cultivation, it will be necessary to add some fresh medium every 3 – 4 days.

## Counter-selection by seed fluorescence using FAST or Sl-FAST2 markers

Selection requires a stereo microscope (with an appropriate magnification) equipped with an RFP filter and UV illumination. There are sure different options for handling seeds for selection – this is just a brief description how we do it:

**Arabidopsis, FAST marker:**

Arabidopsis seeds containing the FAST marker have very strong fluorescence, but are somewhat difficult to handle due to their small size. For rapid selection, we considered it best to spread seeds (not too dense) on a dry piece of printer paper, and to pick desired seeds using a wet toothpick. With some background illumination, this can be done without switching from UV light to white light. Selected seeds are either sown directly, or can be placed on wet Whatman paper for another round of confirmation.

***N. benthamiana*, *Sl*-FAST2 marker:**

*N. benthamiana* seeds containing the *Sl*-FAST2 marker **only demonstrate fluorescence when imbibed** – fluorescence develops within seconds, and is weaker than with FAST in Arabidopsis. We place seeds on a piece of wet Whatman paper for observation. Desired seeds can be picked using appropriate forceps.

## References

Armario Najera, V., Twyman, R.M., Christou, P. and Zhu, C. (2019) Applications of multiplex genome editing in higher plants. Current opinion in biotechnology, 59, 93–102.

Armisen, D., Lecharny, A. and Aubourg, S. (2008) Unique genes in plants: specificities and conserved features throughout evolution. BMC Evol Biol, 8, 280.

Barthel, K., Martin, P., Ordon, J., Erickson, J.L., Gantner, J., Herr, R., Kretschmer, C., Berner, T., Keilwagen, J., Marillonnet, S. and Stuttmann, J. (2020) One-shot generation of duodecuple (12x) mutant Arabidopsis: Highly efficient routine editing in model species. bioRxiv, 2020.2003.2031.018671.

Bensmihen, S., To, A., Lambert, G., Kroj, T., Giraudat, J. and Parcy, F. (2004) Analysis of an activated ABI5 allele using a new selection method for transgenic Arabidopsis seeds. FEBS letters, 561, 127–131.

Boch, J., Bonas, U. and Lahaye, T. (2014) TAL effectors--pathogen strategies and plant resistance engineering. New Phytol, 204, 823–832.

Bolle, C., Huep, G., Kleinbolting, N., Haberer, G., Mayer, K., Leister, D. and Weisshaar, B. (2013) GABI-DUPLO: a collection of double mutants to overcome genetic redundancy in Arabidopsis thaliana. Plant J, 75, 157–171.

Bollier, N., Andrade Buono, R., Jacobs, T.B. and Nowack, M.K. Efficient simultaneous mutagenesis of multiple genes in specific plant tissues by multiplex CRISPR. Plant Biotechnology Journal, **n/a**.

Castel, B., Ngou, P.M., Cevik, V., Redkar, A., Kim, D.S., Yang, Y., Ding, P. and Jones, J.D.G. (2018) Diverse NLR immune receptors activate defence via the RPW8-NLR NRG1. New Phytol.

Cermak, T., Curtin, S.J., Gil-Humanes, J., Cegan, R., Kono, T.J.Y., Konecna, E., Belanto, J.J., Starker, C.G., Mathre, J.W., Greenstein, R.L. and Voytas, D.F. (2017) A multi-purpose toolkit to enable advanced genome engineering in plants. The Plant cell.

Chen, H.W., Bandyopadhyay, S., Shasha, D.E. and Birnbaum, K.D. (2010) Predicting genome-wide redundancy using machine learning. BMC Evol Biol, 10, 357.

Cong, L., Ran, F.A., Cox, D., Lin, S., Barretto, R., Habib, N., Hsu, P.D., Wu, X., Jiang, W., Marraffini, L.A. and Zhang, F. (2013) Multiplex genome engineering using CRISPR/Cas systems. Science (New York, N.Y, 339, 819–823.

Cutler, S. and McCourt, P. (2005) Dude, where’s my phenotype? Dealing with redundancy in signaling networks. Plant physiology, 138, 558–559.

Dang, Y., Jia, G., Choi, J., Ma, H., Anaya, E., Ye, C., Shankar, P. and Wu, H. (2015) Optimizing sgRNA structure to improve CRISPR-Cas9 knockout efficiency. Genome biology, 16, 280.

Davis, A.M., Hall, A., Millar, A.J., Darrah, C. and Davis, S.J. (2009) Protocol: Streamlined sub-protocols for floral-dip transformation and selection of transformants in Arabidopsis thaliana. Plant methods, 5, 3.

Debener, T., Lehnackers, H., Arnold, M. and Dangl, J.L. (1991) Identification and molecular mapping of a single *Arabidopsis thaliana* locus determining resistance to a phytopathogenic *Pseudomonas syringae* isolate. The Plant Journal, 1, 289–302.

Diamos, A.G. and Mason, H.S. (2018) Chimeric 3’ flanking regions strongly enhance gene expression in plants. Plant Biotechnol J.

Ding, T., Huang, C., Liang, Z., Ma, X., Wang, N. and Huo, Y.-X. (2019) Reversed paired-gRNA plasmid cloning strategy for efficient genome editing in <em>Escherichia coli</em>. bioRxiv, 839555.

Edgar, R.C. (2010) Search and clustering orders of magnitude faster than BLAST. Bioinformatics, 26, 2460–2461.

Engler, C., Kandzia, R. and Marillonnet, S. (2008) A one pot, one step, precision cloning method with high throughput capability. PLoS ONE, 3, e3647.

Engler, C., Youles, M., Gruetzner, R., Ehnert, T.M., Werner, S., Jones, J.D., Patron, N.J. and Marillonnet, S. (2014) A Golden Gate Modular Cloning Toolbox for Plants. ACS synthetic biology.

Gantner, J., Ordon, J., Ilse, T., Kretschmer, C., Gruetzner, R., Lofke, C., Dagdas, Y., Burstenbinder, K., Marillonnet, S. and Stuttmann, J. (2018) Peripheral infrastructure vectors and an extended set of plant parts for the Modular Cloning system. PLoS ONE, 13, e0197185.

Gantner, J., Ordon, J., Kretschmer, C., Guerois, R. and Stuttmann, J. (2019) An EDS1-SAG101 Complex Is Essential for TNL-Mediated Immunity in Nicotiana benthamiana. The Plant cell, 31, 2456–2474.

Gao, Y. and Zhao, Y. (2014) Self-processing of ribozyme-flanked RNAs into guide RNAs in vitro and in vivo for CRISPR-mediated genome editing. J Integr Plant Biol, 56, 343–349.

Gelvin, S.B. (2003) Agrobacterium-mediated plant transformation: the biology behind the “gene-jockeying” tool. Microbiol Mol Biol Rev, 67, 16–37, table of contents.

Grützner, R., Martin, P., Horn, C., Mortensen, S., Cram, E.J., Lee-Parsons, C.W.T., Stuttmann, J. and Marillonnet, S. (2020) High-Efficiency Genome Editing in Plants Mediated by A Cas9 Gene Containing Multiple Introns. Plant Communications, 100135.

Hahn, F., Korolev, A., Loures, L.S. and Nekrasov, V. (2019) A modular cloning toolkit for genome editing in plants. bioRxiv, 738021.

He, Y., Zhu, M., Wang, L., Wu, J., Wang, Q., Wang, R. and Zhao, Y. (2018) Programmed Self-Elimination of the CRISPR/Cas9 Construct Greatly Accelerates the Isolation of Edited and Transgene-Free Rice Plants. Mol Plant, 11, 1210–1213.

Jansing, J., Sack, M., Augustine, S.M., Fischer, R. and Bortesi, L. (2019) CRISPR/Cas9-mediated knockout of six glycosyltransferase genes inNicotiana benthamianafor the production of recombinant proteins lacking β-1,2-xylose and core α-1,3-fucose. Plant Biotechnology Journal, 17, 350–361.

Jinek, M., Chylinski, K., Fonfara, I., Hauer, M., Doudna, J.A. and Charpentier, E. (2012) A programmable dual-RNA-guided DNA endonuclease in adaptive bacterial immunity. Science (New York, N.Y, 337, 816–821.

Kannan, B., Jung, J.H., Moxley, G.W., Lee, S.M. and Altpeter, F. (2018) TALEN-mediated targeted mutagenesis of more than 100 COMT copies/alleles in highly polyploid sugarcane improves saccharification efficiency without compromising biomass yield. Plant Biotechnol J, 16, 856–866.

Kroj, T., Savino, G., Valon, C., Giraudat, J. and Parcy, F. (2003) Regulation of storage protein gene expression in Arabidopsis. Development (Cambridge, England), 130, 6065–6073.

Labun, K., Montague, T.G., Gagnon, J.A., Thyme, S.B. and Valen, E. (2016) CHOPCHOP v2: a web tool for the next generation of CRISPR genome engineering. Nucleic Acids Res, 44, W272–276.

Leonhardt, N., Divol, F., Chiarenza, S., Deschamps, S., Renaud, J., Giacalone, C., Nussaume, L., Berthome, R. and Peret, B. (2020) Tissue-specific inactivation by cytosine deaminase/uracil phosphoribosyl transferase as a tool to study plant biology. Plant J, 101, 731–741.

Li, H. and Durbin, R. (2010) Fast and accurate long-read alignment with Burrows-Wheeler transform. Bioinformatics, 26, 589–595.

Li, H., Handsaker, B., Wysoker, A., Fennell, T., Ruan, J., Homer, N., Marth, G., Abecasis, G., Durbin, R. and Genome Project Data Processing, S. (2009) The Sequence Alignment/Map format and SAMtools. Bioinformatics, 25, 2078–2079.

Li, J., Stoddard, T.J., Demorest, Z.L., Lavoie, P.O., Luo, S., Clasen, B.M., Cedrone, F., Ray, E.E., Coffman, A.P., Daulhac, A., Yabandith, A., Retterath, A.J., Mathis, L., Voytas, D.F., D’Aoust, M.A. and Zhang, F. (2016) Multiplexed, targeted gene editing in Nicotiana benthamiana for glyco-engineering and monoclonal antibody production. Plant Biotechnol J, 14, 533–542.

Li, R., Vavrik, C. and Danna, C.H. (2020) Proxies of CRISPR/Cas9 Activity To Aid in the Identification of Mutagenized Arabidopsis Plants. G3 (Bethesda), 10, 2033–2042.

Liu, H., Ding, Y., Zhou, Y., Jin, W., Xie, K. and Chen, L.L. (2017) CRISPR-P 2.0: An Improved CRISPR-Cas9 Tool for Genome Editing in Plants. Mol Plant, 10, 530–532.

Logemann, E., Birkenbihl, R.P., Ulker, B. and Somssich, I.E. (2006) An improved method for preparing Agrobacterium cells that simplifies the Arabidopsis transformation protocol. Plant methods, 2, 16.

Longley, D.B., Harkin, D.P. and Johnston, P.G. (2003) 5-fluorouracil: mechanisms of action and clinical strategies. Nat Rev Cancer, 3, 330–338.

Lowder, L.G., Zhang, D., Baltes, N.J., Paul, J.W., 3rd, Tang, X., Zheng, X., Voytas, D.F., Hsieh, T.F., Zhang, Y. and Qi, Y. (2015) A CRISPR/Cas9 Toolbox for Multiplexed Plant Genome Editing and Transcriptional Regulation. Plant physiology, 169, 971–985.

Ma, X., Zhang, Q., Zhu, Q., Liu, W., Chen, Y., Qiu, R., Wang, B., Yang, Z., Li, H., Lin, Y., Xie, Y., Shen, R., Chen, S., Wang, Z., Chen, Y., Guo, J., Chen, L., Zhao, X., Dong, Z. and Liu, Y.G. (2015) A Robust CRISPR/Cas9 System for Convenient, High-Efficiency Multiplex Genome Editing in Monocot and Dicot Plants. Mol Plant, 8, 1274–1284.

Mali, P., Yang, L., Esvelt, K.M., Aach, J., Guell, M., DiCarlo, J.E., Norville, J.E. and Church, G.M. (2013) RNA-guided human genome engineering via Cas9. Science (New York, N.Y, 339, 823–826.

Mao, Y., Zhang, Z., Feng, Z., Wei, P., Zhang, H., Botella, J.R. and Zhu, J.K. (2016) Development of germ-line-specific CRISPR-Cas9 systems to improve the production of heritable gene modifications in Arabidopsis. Plant Biotechnol J, 14, 519–532.

Minkenberg, B., Xie, K. and Yang, Y. (2017) Discovery of rice essential genes by characterizing a CRISPR-edited mutation of closely related rice MAP kinase genes. Plant J, 89, 636–648.

Najm, F.J., Strand, C., Donovan, K.F., Hegde, M., Sanson, K.R., Vaimberg, E.W., Sullender, M.E., Hartenian, E., Kalani, Z., Fusi, N., Listgarten, J., Younger, S.T., Bernstein, B.E., Root, D.E. and Doench, J.G. (2018) Orthologous CRISPR-Cas9 enzymes for combinatorial genetic screens. Nature biotechnology, 36, 179–189.

Nissim, L., Perli, S.D., Fridkin, A., Perez-Pinera, P. and Lu, T.K. (2014) Multiplexed and programmable regulation of gene networks with an integrated RNA and CRISPR/Cas toolkit in human cells. Molecular cell, 54, 698–710.

Oppenheimer, D.G., Herman, P.L., Sivakumaran, S., Esch, J. and Marks, M.D. (1991) A myb gene required for leaf trichome differentiation in Arabidopsis is expressed in stipules. Cell, 67, 483–493.

Ordon, J., Bressan, M., Kretschmer, C., Dall’Osto, L., Marillonnet, S., Bassi, R., Stuttmann, J.J.F. and Genomics, I. (2019) Optimized Cas9 expression systems for highly efficient Arabidopsis genome editing facilitate isolation of complex alleles in a single generation.

Ordon, J., Gantner, J., Kemna, J., Schwalgun, L., Reschke, M., Streubel, J., Boch, J. and Stuttmann, J. (2017) Generation of chromosomal deletions in dicotyledonous plants employing a user-friendly genome editing toolkit. Plant J, 89, 155–168.

Perera, R.J., Linard, C.G. and Signer, E.R. (1993) Cytosine deaminase as a negative selective marker for Arabidopsis. Plant molecular biology, 23, 793–799.

Pryor, J.M., Potapov, V., Kucera, R.B., Bilotti, K., Cantor, E.J. and Lohman, G.J.S. (2020) Enabling one-pot Golden Gate assemblies of unprecedented complexity using data-optimized assembly design. PLoS ONE, 15, e0238592.

Reis, A.C., Halper, S.M., Vezeau, G.E., Cetnar, D.P., Hossain, A., Clauer, P.R. and Salis, H.M. (2019) Simultaneous repression of multiple bacterial genes using nonrepetitive extra-long sgRNA arrays. Nature biotechnology, 37, 1294–1301.

Römer, P., Hahn, S., Jordan, T., Strauss, T., Bonas, U. and Lahaye, T. (2007) Plant pathogen recognition mediated by promoter activation of the pepper Bs3 resistance gene. Science (New York, N.Y, 318, 645–648.

Shimada, T., Ogawa, Y., Shimada, T. and Hara-Nishimura, I. (2011) A non-destructive screenable marker, OsFAST, for identifying transgenic rice seeds. Plant Signal Behav, 6, 1454–1456.

Shimada, T.L., Shimada, T. and Hara-Nishimura, I. (2010) A rapid and non-destructive screenable marker, FAST, for identifying transformed seeds of Arabidopsis thaliana. Plant J, 61, 519–528.

Sinapidou, E., Williams, K., Nott, L., Bahkt, S., Tor, M., Crute, I., Bittner-Eddy, P. and Beynon, J. (2004) Two TIR:NB:LRR genes are required to specify resistance to Peronospora parasitica isolate Cala2 in Arabidopsis. Plant J, 38, 898–909.

Steynen, Q.J. and Schultz, E.A. (2003) The FORKED genes are essential for distal vein meeting in Arabidopsis. Development (Cambridge, England), 130, 4695–4708.

Stougaard, J. (1993) Substrate-dependent negative selection in plants using a bacterial cytosine deaminase gene. The Plant Journal, 3, 755–761.

Stuitje, A.R., Verbree, E.C., van der Linden, K.H., Mietkiewska, E.M., Nap, J.P. and Kneppers, T.J. (2003) Seed-expressed fluorescent proteins as versatile tools for easy (co)transformation and high-throughput functional genomics in Arabidopsis. Plant Biotechnol J, 1, 301–309.

Stuttmann, J., Hubberten, H.M., Rietz, S., Kaur, J., Muskett, P., Guerois, R., Bednarek, P., Hoefgen, R. and Parker, J.E. (2011) Perturbation of Arabidopsis amino acid metabolism causes incompatibility with the adapted biotrophic pathogen Hyaloperonospora arabidopsidis. The Plant cell, 23, 2788–2803.

Symeonidi, E., Regalado, J., Schwab, R. and Weigel, D. (2020) CRISPR-finder: A high throughput and cost effective method for identifying successfully edited <em>A. thaliana</em> individuals. bioRxiv, 2020.2006.2025.171538.

Thomas, W.J., Thireault, C.A., Kimbrel, J.A. and Chang, J.H. (2009) Recombineering and stable integration of the Pseudomonas syringae pv. syringae 61 hrp/hrc cluster into the genome of the soil bacterium Pseudomonas fluorescens Pf0-1. Plant Journal, 60, 919–928.

Tiraby, M., Cazaux, C., Baron, M., Drocourt, D., Reynes, J.P. and Tiraby, G. (1998) Concomitant expression of E. coli cytosine deaminase and uracil phosphoribosyltransferase improves the cytotoxicity of 5-fluorocytosine. FEMS Microbiol Lett, 167, 41–49.

Torii, K.U., Mitsukawa, N., Oosumi, T., Matsuura, Y., Yokoyama, R., Whittier, R.F. and Komeda, Y. (1996) The *Arabidopsis ERECTA* gene encodes a putative receptor protein kinase with extracellular leucine-rich repeats. The Plant cell, 8, 735–746.

Tsutsui, H. and Higashiyama, T. (2017) pKAMA-ITACHI Vectors for Highly Efficient CRISPR/Cas9-Mediated Gene Knockout in Arabidopsis thaliana. Plant Cell Physiol, 58, 46–56.

van der Biezen, E.A., Freddie, C.T., Kahn, K., Parker, J.E. and Jones, J.D.G. (2002) Arabidopsis *RPP4* is a member of the *RPP5* multigene family of TIR-NB-LRR genes and confers downy mildew resistance through multiple signalling components. Plant Journal, 29, 439–451.

Vidigal, J.A. and Ventura, A. (2015) Rapid and efficient one-step generation of paired gRNA CRISPR-Cas9 libraries. Nat Commun, 6, 8083.

Wang, Z.P., Xing, H.L., Dong, L., Zhang, H.Y., Han, C.Y., Wang, X.C. and Chen, Q.J. (2015) Egg cell-specific promoter-controlled CRISPR/Cas9 efficiently generates homozygous mutants for multiple target genes in Arabidopsis in a single generation. Genome biology, 16, 144.

Warren, R.F., Henk, A., Mowery, P., Holub, E. and Innes, R.W. (1998) A mutation within the leucine-rich repeat domain of the Arabidopsis disease resistance gene RPS5 partially suppresses multiple bacterial and downy mildew resistance genes. The Plant cell, 10, 1439–1452.

Weber, E., Engler, C., Gruetzner, R., Werner, S. and Marillonnet, S. (2011) A modular cloning system for standardized assembly of multigene constructs. PLoS ONE, 6, e16765.

Weiss, T., Wang, C., Kang, X., Zhao, H., Elena Gamo, M., Starker, C.G., Crisp, P.A., Zhou, P., Springer, N.M., Voytas, D.F. and Zhang, F. (2020) Optimization of multiplexed CRISPR/Cas9 system for highly efficient genome editing in Setaria viridis. Plant J, 104, 828–838.

Wolter, F., Klemm, J. and Puchta, H. (2018) Efficient in planta gene targeting in Arabidopsis using egg cell-specific expression of the Cas9 nuclease of Staphylococcus aureus. Plant J, 94, 735–746.

Wu, R., Lucke, M., Jang, Y.T., Zhu, W., Symeonidi, E., Wang, C., Fitz, J., Xi, W., Schwab, R. and Weigel, D. (2018) An efficient CRISPR vector toolbox for engineering large deletions in Arabidopsis thaliana. Plant methods, 14, 65.

Xie, K., Minkenberg, B. and Yang, Y. (2015) Boosting CRISPR/Cas9 multiplex editing capability with the endogenous tRNA-processing system. Proceedings of the National Academy of Sciences of the United States of America, 112, 3570–3575.

Xing, H.L., Dong, L., Wang, Z.P., Zhang, H.Y., Han, C.Y., Liu, B., Wang, X.C. and Chen, Q.J. (2014) A CRISPR/Cas9 toolkit for multiplex genome editing in plants. BMC plant biology, 14, 327.

Zhang, Z., Mao, Y., Ha, S., Liu, W., Botella, J.R. and Zhu, J.K. (2015) A multiplex CRISPR/Cas9 platform for fast and efficient editing of multiple genes in Arabidopsis. Plant cell reports.

Zsogon, A., Cermak, T., Naves, E.R., Notini, M.M., Edel, K.H., Weinl, S., Freschi, L., Voytas, D.F., Kudla, J. and Peres, L.E.P. (2018) De novo domestication of wild tomato using genome editing. Nature biotechnology.

